# Multiparametric Profiling of Single Nanoscale Extracellular Vesicles by Combined Atomic Force and Fluorescence Microscopy: Correlation and Heterogeneity in Their Molecular and Biophysical Features

**DOI:** 10.1101/2021.01.22.427773

**Authors:** Sara Cavallaro, Federico Pevere, Fredrik Stridfeldt, André Görgens, Carolina Paba, Siddharth S. Sahu, Doste R. Mamand, Dhanu Gupta, Samir El Andaloussi, Jan Linnros, Apurba Dev

## Abstract

Being a key player in intercellular communications, nanoscale extracellular vesicles (EVs) offer unique opportunities for both diagnostics and therapeutics. However, their cellular origin and functional identity remain elusive due to the high heterogeneity in their molecular and physical features. Here, for the first time, multiple EV parameters involving membrane protein composition, size and mechanical properties on single small EVs (sEVs) are simultaneously studied by combined fluorescence and atomic force microscopy. Furthermore, their correlation and heterogeneity in different cellular sources are investigated. The study, performed on sEVs derived from Human Embryonic Kidney 293, Cord Blood Mesenchymal Stromal and Human Acute Monocytic Leukemia cell lines, identifies both common and cell line-specific sEV subpopulations bearing distinct distributions of the common tetraspanins (CD9, CD63 and CD81) and biophysical properties. Although the tetraspanin abundances of individual sEVs are independent of their sizes, the expression levels of CD9 and CD63 are strongly correlated. A sEV population co-expressing all the three tetraspanins in relatively high abundance, however, having on average diameters <100 nm and relatively low Young moduli, is also found in all cell lines. Such a multiparametric approach is expected to provide new insights regarding EV biology and functions, potentially deciphering unsolved questions in this field.

## 1. Introduction

Extracellular vesicles (EVs), capable of transmitting biologically active macromolecules, are emerging as key players in intercellular communication. Thus, they have the potential to be used both as non-invasive diagnostic markers as well as therapeutic agents.^[1,2]^ EVs are a heterogenous group of lipid-bilayer nanovesicles (30-2000 nm in diameter) that are secreted by almost all cells and released into the extracellular space.^[3]^ They are broadly divided into three main categories: exosomes (30-150 nm), originating from the endolysosomal pathway, microvesicles (MVs, 50-1000 nm), formed by direct outward budding of the plasma membrane, and apoptotic bodies (500-2000 nm), derived from apoptotic cells.^[4]^ However, in recent years it has been increasingly understood that cells secrete subpopulations of EVs that make them functionally more diverse.^[5–8]^ EVs can impact neighboring cells or cells at a distance, as they contain lipids, proteins, or nucleic acid species from the source cell, and have the unique ability to convey these macromolecules via a highly advanced system of intercellular communication. Consequently, they play an important role in numerous physiological and pathophysiological processes, including immune regulation and cancer development.^[4,9]^ Different cell types may produce distinct repertoires of vesicles, reflecting the physiological state of the cell.^[10]^ Understanding the EV biology and physiological relevance, therefore, requires the capacity to resolve their phenotypic variations and identify their functional relations. However, the high heterogeneity of EVs even from a single cell type renders analyses by most available technologies rather ineffective, as they are predominantly based on average properties.^[11]^ Therefore, there has been an increasing effort for analysis of EVs in general and small EVs (sEVs)/exosomes in particular at a single particle level.^[12–18]^ On the contrary, the analysis of single sEVs on a sufficiently large and representative population is also extremely challenging, as their size lies well below the optical resolution and they are very weak light scatterers.^[19]^ In addition, many sEV properties, such as the abundance and types of surface proteins, their sizes, genetic cargo, mechanical properties etc., all seem to play important roles in EV functions,^[13,20]^ indicating the need to include a number of these parameters to identify subtypes within a given sEV population. Most of the reported single vesicle studies so far have only been able to identify sEV subpopulations based on their protein expressions, investigated mainly by immunofluorescent staining approaches.^[12,15,18]^ Although these studies reveal various interesting features of distinct EV subtypes, they may not be sufficient to fully address the issue of heterogeneity. Going a step ahead, recently, Tian et al. successfully combined immunofluorescent staining with side scattering technique to simultaneously measure both the size and protein expression of single EVs, identifying previously unknown EV subtypes.^[13]^ Daaboul et al. also achieved similar results by combining antibody-based capture with interferometric imaging.^[14]^ Recent studies investigating the EV mechanical and structural properties have also found various EV subpopulations bearing distinct mechanical behavior,^[21,22]^ which might be influenced by their parent cell and/or their pathophysiological state, thereby, resulting in variations in their interactions with cells.^[21,23]^ These observations obviously raise new questions, as to which degree are these properties correlated? If not, then what relation do they carry with their parent cells?

Although intriguing, such a study would require a platform that can accurately determine the size, membrane protein composition and mechanical properties of sEVs, in their physiological environment. However, these properties cannot be simultaneously measured by the available technologies, such as Nanoparticle Tracking Analysis (NTA), Interferometric Reflectance Imaging (IRI), Resistive Pulse Sensing (RPS) etc. Moreover, the size estimation performed by such techniques is limited in accuracy and/or detection range.^[24]^ By combining fluorescence (FL) and atomic force microscopy (AFM), we performed, for the first time, such a multiparametric analysis on single sEVs derived from three different cell lines, namely Human Embryonic Kidney 293 (HEK293), Cord Blood Mesenchymal Stromal (cbMSC) and Human Acute Monocytic Leukemia (THP1), respectively. In our study, the AFM, installed on top of the fluorescence microscope and aligned with the optical axis, allowed accurate combined measurements on individual sEVs, thanks to a very precise spot identification approach. To minimize size/shape alteration and damage of the soft nanovesicles, force curve-based imaging (or quantitative imaging, QI) mode was performed in liquid environment.^[25]^ Aiming to delineate the sEV heterogeneity manifested in their molecular, morphological and mechanical properties, we investigated the abundance of common tetraspanins, e.g. CD9, CD63 and CD81 by immunostaining approach and combined that with high resolution AFM analysis of their sizes and stiffnesses (Young Modulus, YM). The results confirmed the high heterogeneity of the sEVs in their physical and molecular properties, highlighting the presence of sEV subpopulations with possible different functions/biogenesis routes. Overall, our data revealed the presence of vesicles with a large range of diameters (30-200 nm), Young moduli (~0.1-25 MPa) along with a large variation in the distribution of the investigated membrane proteins. While sEVs from all the cell lines displayed a correlation between the expression levels of CD9 and CD63, the abundance of the investigated tetraspanins were found to be independent of the sEV diameter. In addition, some common and cell-line specific features were also observed in their molecular, morphological and mechanical properties, which highlights the prospect of the proposed method for highly accurate and multiparametric EV analysis.

## 2. Results

### 2.1 Single EV technology

**Figure 1** schematically demonstrates our measurement platform. sEVs were captured and conjugated to a glass coverslip by using covalent coupling via glutaraldehyde(GA)-amine interaction (Figure 1A, see Experimental Methods for details).^[26]^ This was necessary to ensure stability of the sEVs during the entire investigation period, as needed for AFM imaging as well as for immunostaining. Moreover, it assures that the sEV capture protocol is efficient, i.e. capable of capturing a significant population from the sample volume and is unbiased to size and expression level. We first tested and optimized the method with HEK293 cell line-derived sEVs, engineered to overexpress CD63 tagged with m-neongreen (mNG) protein.^[27]^ For these sEVs, referred as mNG-EVs hereafter (Figure 1A, top), the sEV-conjugation step was followed by PBS washing and combined FL-AFM imaging. For the wild type sEVs (HEK293, cbMSC and THP1) used in the study, referred as wt-EVs hereafter (Figure 1A, bottom), this sEV conjugation step was followed by, i) surface blocking using tris-ethanolamine (Tris-ETHA) and casein, to minimize non-specific binding (NSB) of the antibodies, ii) fluorescently tagged antibody incubation, and iii) washing steps prior to imaging (see Experimental Methods for details).

**Figure 1.**
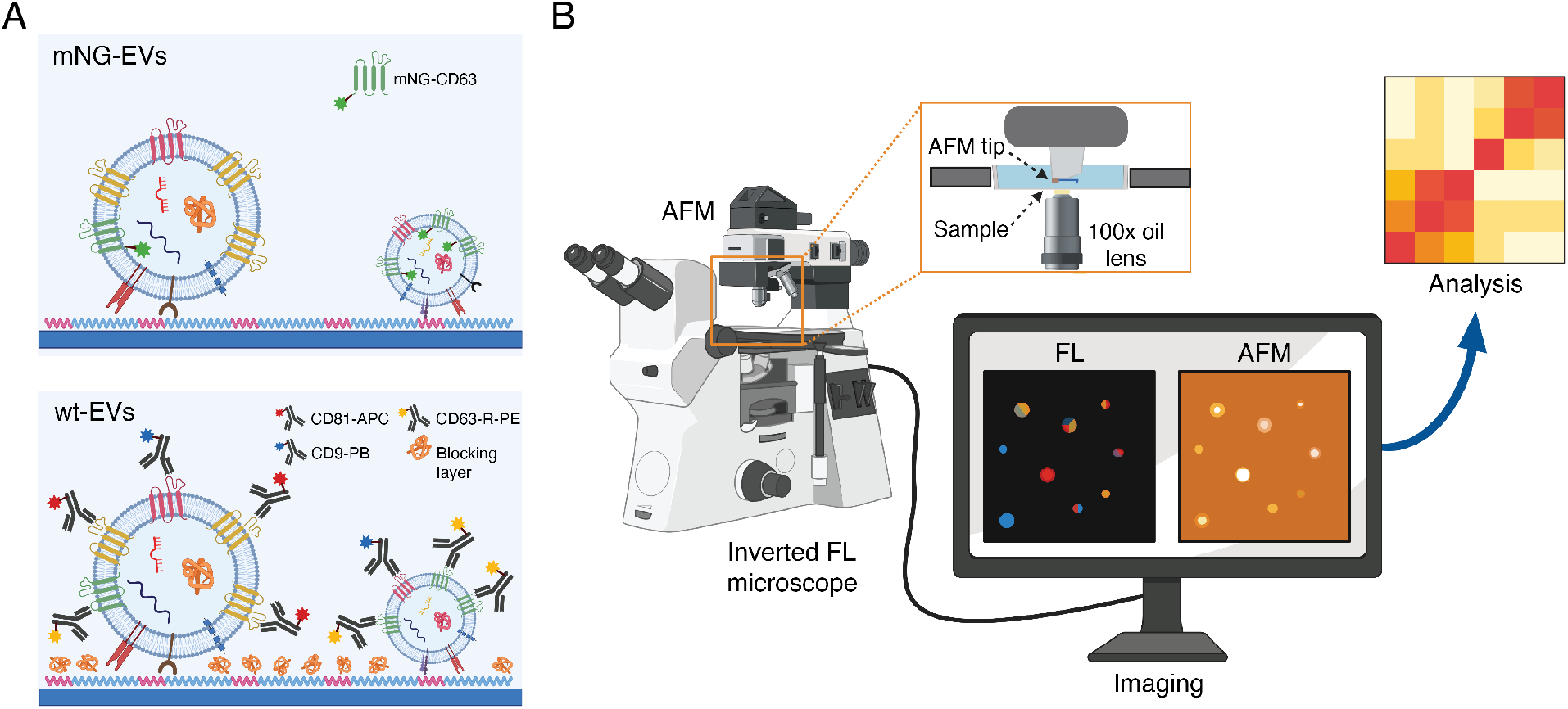
Schematic of the single EV platform. (A) Schematic of the sEV capture procedure for both engineered m-neongreen tagged sEVs (mNG-EVs) and wild type sEVs (wt-EVs). While the mNG-EVs had the fluorescent mNG-CD63 construct, the wt-EVs were targeted using three different fluorescently labelled antibodies. (B) Schematic of the imaging platform, including inverted FL microscope below the sample support with the EV substrate (inset) and AFM above it, and aligned with optical axis. The FL and AFM images were visualized on a screen and then processed and analyzed using different software. This figure was realized using BioRender.com.

On these wt-EVs, we analyzed the distribution of the widely abundant tetraspanins CD9, CD63, and CD81.^[18]^ It is well-known that the size of the antibodies (~5-10 nm^[28]^), in addition to various degree of abundance of their respective membrane proteins, may produce significant steric hindrance. This is particularly true when the antibodies are conjugated in sequence,^[29]^ leading to a notable error in determining the relative membrane protein levels. In order to minimize such error, we prepared a solution containing all the three fluorescently labelled antibodies recognizing these markers, and let it incubate with the sEVs for 2 h. Particularly, the solution included anti-CD81-Allophycocyanin (CD81-APC, 2nM) antibody, anti-CD63-R-Phycoerythrin (CD63-R-PE, 2nM) antibody and anti-CD9-PacificBlue (CD9-PB, 2 nM) antibody in 1x PBS (see Experimental Methods for details). The selected fluorophores could be imaged simultaneously, showing negligible crosstalk (Figure S1). Both the mNG-EVs and wt-EVs were isolated using ultrafiltration (UF) and bind-elute size exclusion chromatography (BE-SEC).^[30]^ The isolated vesicles were characterized by nanoparticle tracking analysis (NTA, Malvern system) and Western Blot (WB, for mNG-EVs)/multiplex bead-based flow cytometry assay (MACSPlex Exosome Kit, for wt-EVs, see Experimental Methods for details). Figure S2 shows the results of such sEV characterization. After the antibody incubation and washing steps, we performed combined FL-AFM imaging, using the setup illustrated in Figure 1B. The FL images were used to qualitatively estimate the abundance of the analyzed tetraspanins on individual sEVs, while the AFM scans were used to estimate their sizes (equivalent diameters) and mechanical properties (Young Moduli). The measurements were performed in liquid (1x PBS) by inserting the substrate in a transparent well (Figure 1B, inset), which was placed in between the objective lens and AFM tip for the combined FL-AFM approach.

### 2.2 Platform characterization on engineered mNG-EVs

As mentioned before, we first optimized and characterized the platform with mNG-EVs to ensure that, i) the captured EVs remained stable during the entire investigation period, including washing steps and measurements, ii) the EV capture protocol was efficient and unbiased towards EV size and protein expression, iii) the platform had sufficiently large dynamic ranges for size and surface expression measurements, iv) the approach discriminated single particles. **Figure 2**A illustrates a representative FL image of covalently captured mNG-EVs, showing distinct bright spots that could be assigned to single particles. A control measurement performed on an identical substrate but without any EVs clearly showed no such detectable luminescent spots (Figure S3). In order to investigate the stability of the captured EVs and emulate the conditions necessary for antibody staining, we assessed the effect of multiple washing steps on the retention of the EVs on the substrate. The measurements revealed that the vesicles remained immobilized even after multiple washings with PBS or water, with a particle variation <1% after the third washing step (data not shown). Furthermore, the vesicles remained attached to the substrate for at least 12 h, indicating that the covalent capture ensured sufficient EV stability. To evaluate the capture efficiency of our protocol, we counted the number of fluorescence spots for different sEV concentrations and compared the results with the unitary slope derived from NTA. For the analysis, we first optimized the surface conjugation protocol to ensure that almost all the EVs from the solution interacted and attached to the substrate homogeneously and with minimum losses. Considering homogeneous immobilization, we then converted the area density of particles to the initial volume concentration. As Figure S4 shows, the number of captured sEVs was 3 times lower than the nominal particle counts obtained by NTA, indicating a capture efficiency of 33% only. However, given that NTA also detects non-EV particles^[31]^, that not all the EVs expressed the mNG-CD63 construct and some weakly luminescent EVs may exist below the detection limit, we expected our capture protocol to be far more efficient than 33% (Figure S4). For protein expression analysis, we considered the integrated FL intensity distribution (Figure 2B), obtained after background subtraction (Zeiss Data Analysis software, see Experimental Methods for details). The data showed an exponentially decreasing FL distribution ranging from ~680 a.u. to ~5×10^5^ a.u., with increasing number density towards the low intensity values.

**Figure 2.**
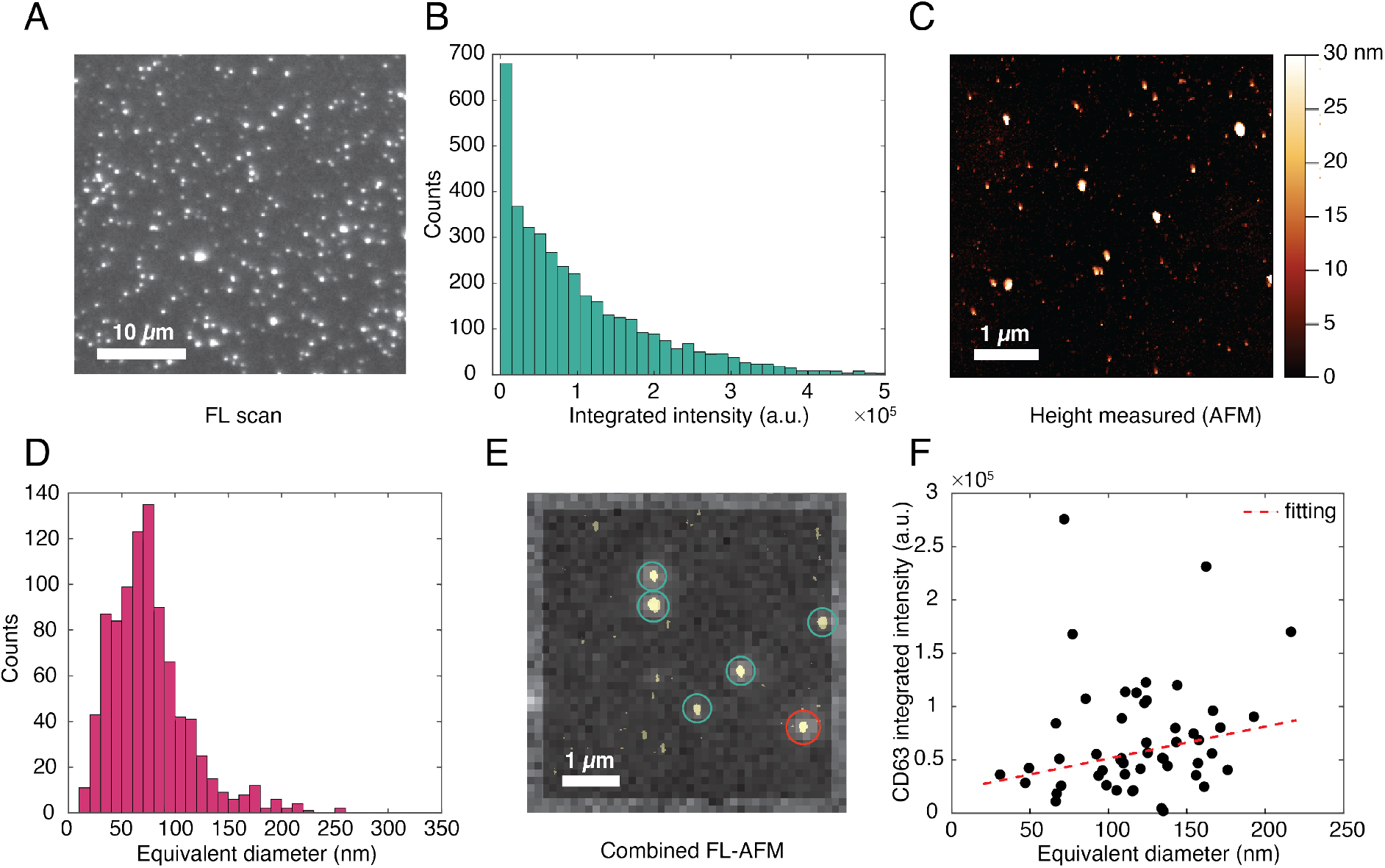
Platform characterization and optimization with mNG-EVs. (A) Representative FL image of mNG-EVs captured on a glass substrate. (B) FL integrated intensity distribution of the captured mNG-EVs. (C) Representative AFM scan in liquid of mNG-EVs captured on a separately prepared glass substrate. Height threshold for EV identification was set at 15 nm. (D) AFM equivalent diameter distribution of the captured mNG-EVs (height > 15 nm), obtained assuming surface area conservation. Distribution obtained from 17 scans of 5 *μ*m × 5*μ*m (total area=425 *μ*m^2^) (E) Representative image of the combined FL (outer image with diffuse encircled spots) and AFM (inner image with yellow solid particles) experiments. (F) Correlation plot between the size (equivalent diameter, x-axis) and the fluorescence intensity (integrated intensity, y-axis) of single vesicles. The fitted curve was obtained by Bi-square linear fitting.

Next, we optimized the AFM parameters by separately measuring sEVs in liquid using soft tips specially designed for biological samples (see Experimental Methods for details). Figure 2C and S5 show representative AFM height images obtained from a substrate with mNG-EVs and an identically prepared control substrate without EVs, respectively. The scans show clear differences between the substrates with EVs (Figure 2C) and without (Figure S5) for a particle height higher than 15 nm, which was set as a threshold for sEV identification. Since the vesicles appeared to assume a flattened shape on the substrate, with the diameters being about twice the heights, for further analysis we considered the EV equivalent diameters which were estimated by assuming surface area conservation.^[32]^ The equivalent diameter distribution in Figure 2D suggests that ~25% of the total number of particles had diameters below 50 nm and that the mean particle diameter was ~76 nm. The presence of such small particles could be due to small sized-EVs/exomeres and/or proteins/protein aggregates that often remain undetected by NTA and scattering-based methods.^[24]^ The peak of the distribution appeared at around 70 nm, which is shifted by 40 nm as compared to the size estimation performed by NTA (Figure S2A). Overall, both the FL intensity and AFM diameter distributions showed that our capture protocol was unbiased towards the EV size and protein expression. Furthermore, the sufficiently large dynamic ranges of such distributions (including but not limited to 680-5×10^5^ a.u. for FL intensity and 10-250 nm for AFM equivalent diameter) confirmed the feasibility of our platform for accurate sEV size and surface marker expression analyses.

Once validated and optimized separately, we performed combined FL and AFM measurements on the mNG-EVs. Figure 2E and S6 show representative images from such measurements of different substrate areas, obtained by overlapping the FL images with the AFM scans of the same spots. Since surface roughness, small spatial shifts in AFM scan, etc. can introduce significant uncertainty in the precision of the image overlap, despite having optical axis alignment of the two systems, we etched a series of nano- and microscale patterns on the coverslips prior to functionalization. This step was crucial for precise spot identification. Overall, the AFM data could also be used to identify the FL spots corresponding to a single sEV (Figure 2E, green circles) or groups of multiple particles/EVs (Figure 2E and S6, red circles). The latter were excluded from the single particle analysis. As shown, the results suggested that almost all of the fluorescence spots were generated by single EVs, with some exceptions (~10%) where a spot corresponded to two or more particles. Figure 2F shows the distribution of mNG-CD63 expression level as a function of the sEV size. For the analysis, we only considered the FL spots corresponding to a single AFM particle and related them to the vesicle equivalent diameters. As suggested from the relatively low slope of the fitting curve and R-square coefficient (R-square=0.50), the number of mNG-CD63 proteins was not strongly proportional to the vesicle diameter but was distributed rather stochastically within the expected sEV size range. Moreover, the FL intensity on single vesicles ranged from ~2000 a.u. to ~3×10^5^ a.u. (Figure 2F), with most of the sEVs showing a FL count <1.5×10^5^ a.u.. This indicated that some of the higher FL intensity values in Figure 2B were likely attributed to agglomerated vesicles. Furthermore, the maximum FL intensity was ~100 times larger than the minimum value. This data range is similar to that reported in other studies, which show that EVs can capture from 1 to ~50-60 copies of a protein.^[13]^

### 2.3 Fluorescence analysis on wt-EVs

Once validated, we applied the method to profile wt-EVs from the three different cell lines HEK293, cbMSC and THP1. In particular, we examined the distribution of the widely abundant tetraspanins CD9, CD63, CD81 and analyzed their dependence on the size and Young Modulus of the corresponding sEVs. The choice of these cell lines was justified by their availability to others in the field (standard cell lines), their known differences between the surface protein composition,^[18,27]^ and the absence of such a multiparametric study at a single vesicle level. **Figure 3**A shows a representative FL image of sEVs stained with all the three antibodies. In the image, the presence of the fluorophores corresponding to CD9, CD63 and CD81 are indicated by the colors blue, yellow and red, respectively. Distinct bright spots with sufficiently low background clearly indicated good specificity of the immunostaining protocol.

**Figure 3.**
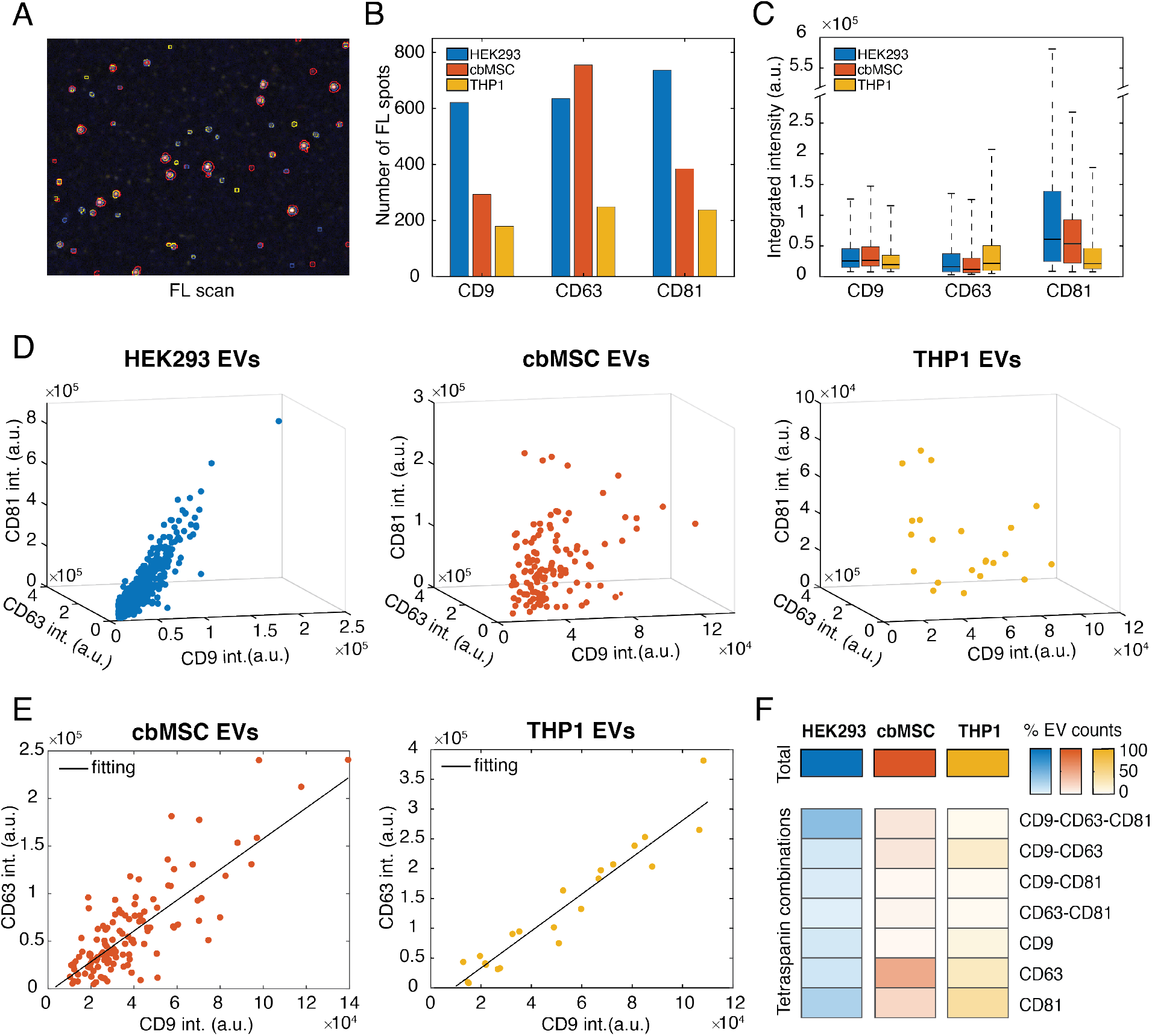
FL analysis on the three wt-EV samples HEK293, cbMSC and THP1. (A) Representative FL image of EVs targeted with CD9-PB, CD63-R-PE and CD81-APC antibodies, showing vesicles co-expressing all markers simultaneously (three colors) and vesicles only expressing two or one markers. (B) Number of positive EVs detected in a microscope area for each tetraspanin analyzed and cell line. (C) Box plots summarizing the integrated intensity distributions for all the markers and samples analyzed. Whisker value set to 4. (D) Correlation of the FL integrated intensities for the CD9-CD63-CD81 positive EVs for the HEK293 (left, blue), cbMSC (middle, red) and THP1 (right, yellow) EVs, respectively. (E) Correlation of the CD9 and CD63 FL intensities for the CD9-CD63-CD81 positive EVs of cbMSC (red) and THP1 cell lines (yellow). (F) Heatmap showing the amount of sEVs present for each protein combination subgroup. For example, CD9-CD63-CD81 label indicates the number of EVs positive for the three markers simultaneously, CD9-CD63 the number of EVs positive only for CD9 and CD63, whereas CD9 the number of EVs only positive for CD9 protein (no CD63, no CD81 detected). Numbers of EVs per subgroup expressed as a percentage of the total number of EVs detected for each cell line (100%).

The same was further verified with control measurements, where the antibodies were incubated on functionalized substrates without sEVs (Figure S7) or where isotype control antibodies were incubated on substrates with sEVs (Figure S7). The FL results presented in Figure 3 also show the differences among the analyzed samples, in terms of both the abundances and distributions of the tetraspanins within the sEV populations. In particular, Figure 3B shows the number of sEVs that were detected to express the different tetraspanins in each of the cell lines, from a microscope area (133.12 *μ*m × 133.12 *μ*m). As the EV capture protocol was unbiased to the membrane proteins, we expected a similar number of vesicles binding to the substrate for all the cell lines. Therefore, Figure 3B could be used to compare the populations of sEVs that were positive to the different tetraspanins in these cell lines. The results (Figure 3B) revealed that for the HEK293 cell line, the number of sEVs positive to each of the tetraspanins were similar, with a slight enrichment in CD81 expressing EVs. The cbMSC sEVs, instead, were more abundant in CD63 positive EVs than the other two.^[33]^ Similar to HEK293, the THP1 sEVs also did not show prevalence of a particular marker (Figure 3B). Figure 3C presents a box plot graphically depicting the distribution of the tetraspanin intensities collected from each of the sEVs for the three cell lines. Being such integrated intensities proportional to the number of tetraspanins on each vesicle, the data showed that the HEK293 EVs had the highest abundance of CD81 per sEV compared to the other two cell lines. The THP1 EVs showed instead the highest levels of CD63 per sEV, while none of the analyzed cell lines showed prevalence of CD9 per vesicle. When analyzing the sEV population expressing the three tetraspanins simultaneously, the results revealed interesting correlations, as presented in Figure 3D. In particular, we observed that, while for the HEK293 sEVs, the increase of one marker, e.g. CD9, was accompanied by a linear increase of the other two markers, i.e. CD63 and CD81 (R-square_tot_=0.84, R-square_CD9-CD63_=0.88, R-square_CD9-CD81_=0.83, R-square_CD63-CD81_=0.81), for the cbMSC and THP1 EVs, there was not a similar correlation among all the three tetraspanins (R-square_tot_=0.2 and R-square_tot_=0.27, respectively). However, the intensities of CD9 and CD63 were correlated for the cbMSC and THP1 sEVs in this group, with both linearly increasing with each other (Figure 3E, R-square=0.72 for cbMSC and R-square=0.92 for THP1 EVs). The same correlation between these two tetraspanins was also observed for the sEV populations only positive to CD9-CD63 for all the three cell lines (Figure S8), indicating that this is a general feature of the cell lines studied here. The data also showed remarkable differences in the distribution of the marker combinations among the samples. This is highlighted in the heatmap in Figure 3F, which illustrates the proportions of different EV sub-populations categorized according to their tetraspanin combinations. As presented, both the HEK293 and cbMSC cell lines showed similar amount of total tetraspanin positive sEVs (992 and 1005, respectively) in the investigated microscope area, while for THP1, the amount stood at nearly 50% (502 tetraspanin positive sEVs detected). Within their respective populations, the fraction that contained all the three tetraspanins were about 40%, 12% and 4%, respectively for HEK293, cbMSC and THP1 EVs. This indicated that the co-presence of all the three tetraspanins was a far more dominant feature of the HEK293 sEVs than the other two. On the other hand, the cbMSC sEVs seemed to dominantly favor vesicles only positive for CD63 (~48% of the population), while the THP1 sEVs showed preference to vesicles only positive for CD81 (~38% of the population). Another common feature, as appears under the group of sEVs co-expressing only two tetraspanins simultaneously (Figure 3F), is that the co-enrichment of CD9-CD63 combination was favored compared to the other two tetraspanins combinations among all the three cell lines. This was particularly dominant in THP1 EVs, where ~20% of the total population displayed this feature.

### 2.4 FL clustering analysis on wt-EVs

In order to understand how the abundance of the various tetraspanins per sEV influence the subpopulations identified in Figure 3F, we mapped the sEV FL intensity data onto a two-dimensional plane using t-Distributed Stochastic Neighbor Embedding (tSNE) method. For the analysis, we first optimized the mappings for each cell line according to perplexity values and convergence (see Experimental Methods for details). The results of the analysis are presented in Figure S9, where each point represents a single vesicle. We emphasize here that the abundance of the tetraspanins was qualitatively compared by taking the FL intensity of each tetraspanin channel normalized by the minimum detectable intensity (weakest spot from a single EV) for the same channel and cell line. This value is proportional to the amount of each type of protein present on each vesicle. As shown in Figure S9, the optimal parameters identified 5 main clusters for the HEK293 EVs, 3 clusters for cbMSC EVs and 5 clusters for the THP1 EVs. **Figure 4**A presents the constituent tetraspanin combinations in each of the identified clusters, while Figure 4B shows the distributions of their FL intensity values. The analysis suggested that for the HEK293 EVs, the most populated cluster (cluster 2, 49% of the sEV population, Figure S9 and 4A) was composed mainly by the vesicles expressing the three tetraspanins simultaneously and by those only positive for CD9-CD63 (Figure 4A). Although the intensities spread over a large range (cluster 2, Figure 4B), on average this group of vesicles showed the highest intensity values for all the tetraspanins, indicating an enrichment of such markers. The second most populated cluster (cluster 4, 26% of the sEV population, Figure S9 and 4A), in this cell line was composed of vesicles either positive for CD81 only or for CD63-CD81. Intensity comparison showed that compared to cluster 2 (Figure 4B), all the other clusters showed low levels of tetraspanin abundance per vesicle.

**Figure 4.**
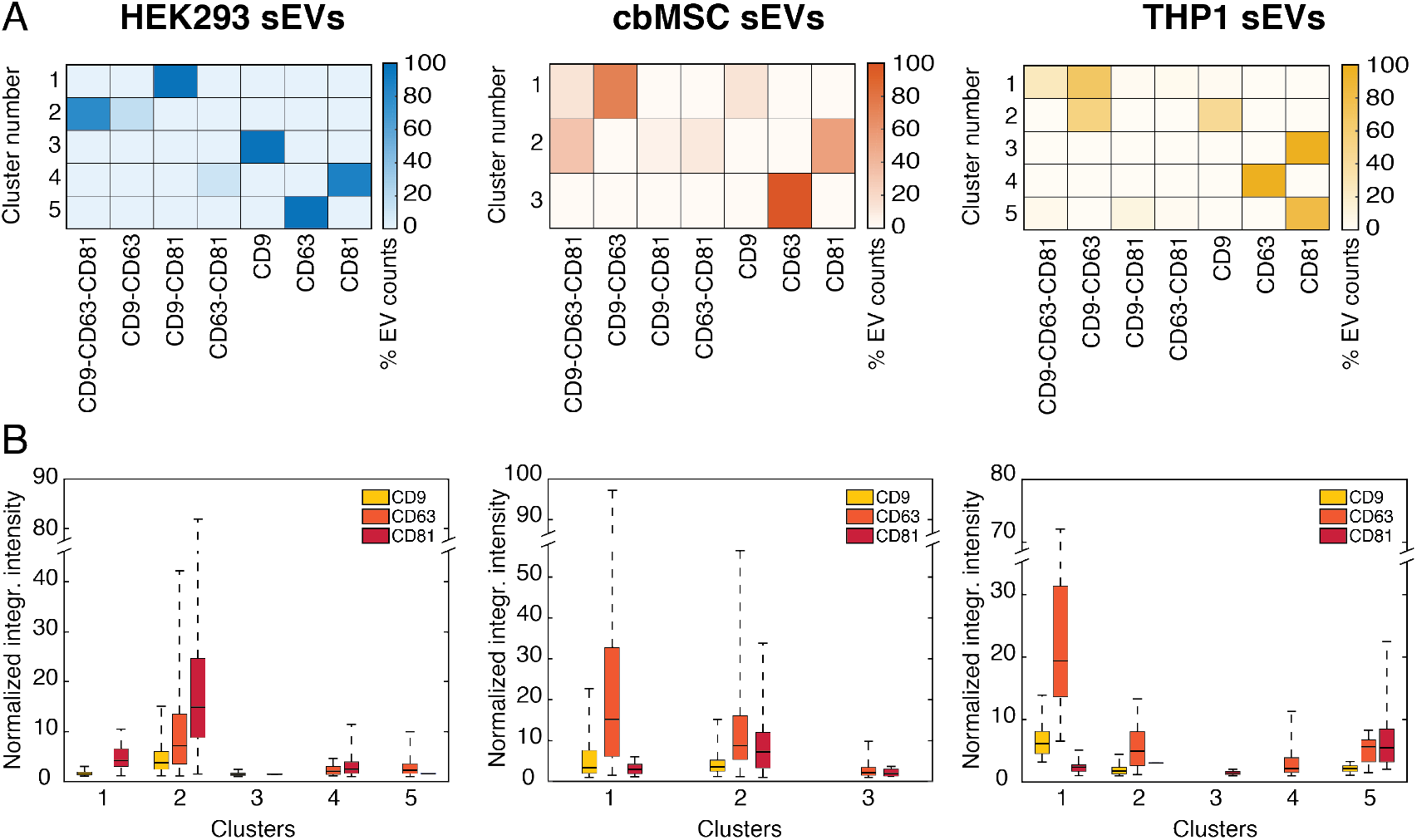
Clustering analysis on the HEK293, cbMSC and THP1 wt-EVs. The results from the different cell lines are separated in three different columns. (A) Heatmaps of the tSNE derived marker expression profiles, showing the tetraspanin combinations within the identified clusters (5 for HEK293 EVs, 3 for cbMSC EVs and 5 for THP1 EVs). (B) Box plots of the normalized FL intensity distributions of the three tetraspanins for the tSNE derived clusters.

The results obtained for the cbMSC sEVs revealed a quite different behavior. In this case, the clustering algorithm identified 3 main clusters. Unlike HEK293, in this cell line the EV subpopulation co-expressing all the three tetraspanins split into two clusters, one of them including also the EVs positive only for CD9-CD63 (cluster 1, 17% of the sEV population, Figure 4A) and the other one including the EVs expressing CD81 (cluster 2, 35% of the sEV population, Figure 4A). Interestingly, the CD9-CD63-CD81 positive vesicles clustering with the CD9-CD63 expressing sEVs (cluster 1) showed the highest levels of CD9 and CD63, while the sEVs clustering with the CD81 expressing vesicles (cluster 2) showed the highest levels of CD81, but intermediate levels of CD9 and CD63. Finally, the data for the THP1 sEVs suggested a similar behavior as the cbMSC vesicles.

### 2.5 Distribution of EV physical parameters within clusters

Following fluorescence analysis and clustering, we analyzed how the vesicle physical properties, e.g. size and YM, related to their protein expressions, in order to further characterize the EV heterogeneity and investigate the role played by these parameters. As reported earlier, other than size profiling, AFM can also accurately determine particle adhesion, stiffness or deformability, which may be responsible for different cellular interactions.^[21]^ **Figure 5** summarizes these results, showing the distribution of the sEV size (Figure 5A) and YM (Figure 5B) within the identified clusters. The YM values were used as a measure of the vesicle stiffness. For this purpose, we considered the vesicles randomly scanned by AFM on different substrate areas of the initially acquired FL area, in order to obtain unbiased measurements (see Experimental Methods for details). Figure S10 and S11 show a representative combined FL-AFM measurement performed on the HEK293 wt-EVs and the size distributions of the analyzed vesicles, respectively. Due to the low throughput of AFM in image acquisition, we could only analyze a fraction of those vesicles identified by fluorescence imaging. This was necessary to ensure the best image resolution. However, the acquired data points were representative of the whole sEV populations (Figure S12 and S13). As presented in Supporting Information, the vesicles analyzed by AFM were uniformly distributed among all the different clusters (Figure S12) and showed similar tetraspanin distributions (Figure S13) as the whole sEV dataset (Figure 4B).

**Figure 5.**
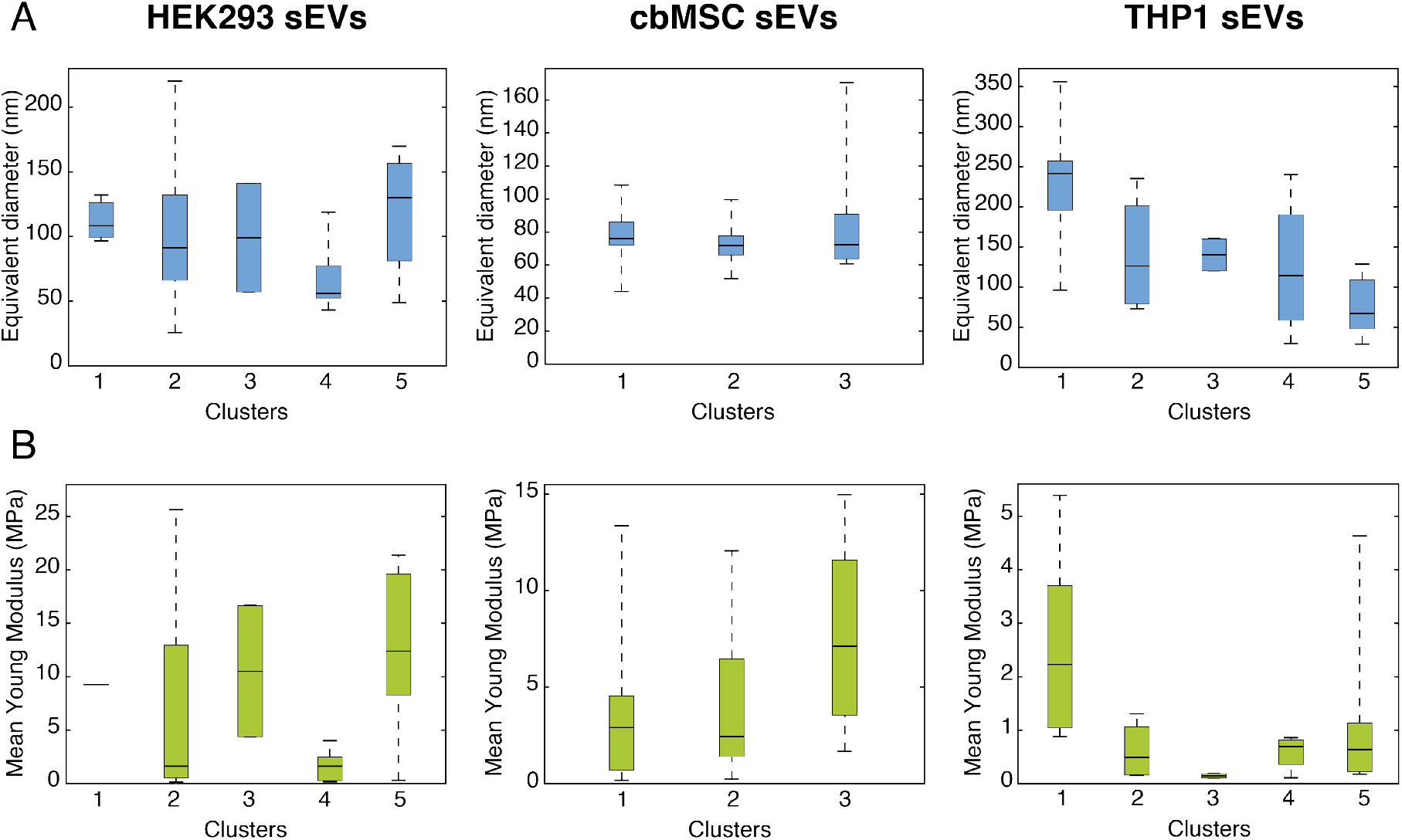
Distribution of (A) size and (B) Young Modulus of the sEVs analyzed by AFM within the previously identified clusters based on FL intensities.

Figure S14 shows the plots of the tetraspanin intensities as a function of sEV equivalent diameters for the three cell lines. Clearly, the vesicle size did not increase with tetraspanin intensity, suggesting that high expressing vesicles did not necessarily have large diameters. However, when analyzing the size distribution within the identified FL clusters (Figure 5A), the data revealed some interesting features. In particular, for the HEK293 EVs, the vesicles expressing the three proteins simultaneously (cluster 2) had on average small diameters (<100 nm), but also the largest variations. The diameters of the sEVs in this cluster were distributed in the range from ~30 nm to ~220 nm, thereby confirming the existence of vesicles with sizes <50 nm in this subpopulation. However, the most intense sEVs in this group were centered immediately around 100 nm in diameter (Figure S14). On the other hand, the sEVs positive for CD81 only or those expressing CD81 in combination with CD63 (cluster 4) had the smallest mean diameters (~60 nm). The remaining clusters (1, 3, 5), containing the sEVs expressing low levels of tetraspanins, showed on average larger diameters. The results for the cbMSC and THP1 sEVs revealed some analogies but also several differences. In both these samples the sEVs expressing the three tetraspanins, which split between two clusters, showed two different behaviors. In particular, those vesicles coupling with the CD9-CD63 expressing sEVs (clusters 1 for both cbMSC and THP1 sEVs) had on average larger diameters (Figure 5A) than those coupling with the CD81 expressing ones (Figure 5A, cluster 2 for cbMSC sEVs and cluster 5 for THP1 sEVs). However, while for the THP1 sEVs, the vesicles expressing the three tetraspanins showed the largest diameter variations, for the cbMSC ones, these vesicles were distributed within a narrow range (mostly around 70-80 nm, Figure S14 and S15). For cbMSC, while most of the analyzed sEVs had diameter <100 nm, the population expressing CD63 only (cluster 3, Figure 5A) showed a large spread in diameter reaching up to 160 nm.

The analysis of the distribution of the YM within the clusters suggested that overall the vesicle stiffness followed a similar trend as the EV equivalent diameters. Despite sample-to-sample variations in stiffness, the YM values for single vesicles ranged from ~170 kPa to ~25 MPa. Although more detailed studies and experiments need to be performed on this aspect, the preliminary results in Figure 5B showed that the sEVs were also widely heterogeneous in their stiffness, with the larger vesicles appearing to be stiffer than the smaller ones. In particular, for all the samples, the small CD81 only positive vesicles were much softer than the other clusters, while the larger CD9-CD63 expressing sEVs were stiffer (data not shown). In a similar trend as the diameter, the stiffest EVs for the cbMSC cell line were the largest CD63-only expressing sEVs (cluster 3, Figure 5B). The comparison of the YM (Figure 5B) with the FL intensities (Figure 4B) revealed that the presence of the tetraspanins seemed to soften the vesicles for the HEK293 and cbMSC samples. Indeed, the sEVs with a higher amount of proteins showed smaller YM on average. The THP1 sEVs, instead, showed an opposite behavior.

## 3. Discussion

Over the past decade, many scientific investigations have highlighted the importance of EVs in intercellular communication and also their role in pathological conditions.^[1,2,4,9]^ EVs, being generated from either cell membrane budding or endosomal invagination, are known to capture cell specific proteins, RNAs or lipids.^[6]^ However, the mechanisms of this cargo packaging remain largely elusive so is the identity of functionally distinct EV subsets. This is because the vesicles carry a large number of molecular and biophysical signatures, which necessitates their individual characterization with an accurate and multiparametric approach to be able to delineate their common and distinct features. In this context the proposed method offers a unique advantage as a number of EV parameters such as their membrane protein composition and abundance, their size and mechanical properties can be investigated with very high resolution and accuracy. As a proof of concept, we developed a sample preparation protocol with mNG-EVs to ensure efficient and unbiased EV capture irrespective of their size (Figure 2D) and protein expression (Figure 2B). The strategy also showed to maintain the stability of the captured vesicles for the investigation. As presented in Figure 2D, S10 and S11, AFM can detect particles as small as 10-20 nm, which are below the reported size range of EVs.^[4]^ Although it is still not known whether these particles are EVs or exomeres,^[34]^ the data highlight the capability of the method to characterize EVs in their small size range (i.e. 30-40 nm, Figure S10) as well as to investigate the purity of the EV isolation methods. However, a major benefit stems from the fact that the combined FL-AFM measurements can also easily discriminate single vesicles from other closely spaced particles or EV groups (Figure 2E) which are difficult to identify by fluorescence and/or scattering based approaches.

The results obtained by analyzing the sEVs from the three different cell lines in general show large heterogeneity in the sEV populations, displaying both common and distinct features among the cell lines. As presented in Figure 3B and 3C, we found that CD81 is a more frequently and abundantly expressed tetraspanin in HEK293 EVs, a feature also reported earlier.^[35]^ In comparison, cbMSC EVs show a prevalence of CD63 among the tested proteins. This is also in agreement with a previous report.^[33]^ Furthermore, HEK293 sEVs are more enriched with the analyzed tetraspanins compared to the other two samples (Figure 3B),^[27]^ while THP1 sEVs show the lowest numbers of tetraspanin positive vesicles. However, in comparison with the previous reports our combined method has the advantage to complement the protein abundance data with the vesicle size and stiffness (YM), thereby further deciphering EV heterogeneity. For all the cell lines included here, the results also suggested the presence of a co-localization and correlation between CD9 and CD63 proteins, with both increasing (or decreasing) with each other on single vesicles. Moreover, for all the cell lines, we detected a sEV subpopulation which was composed of vesicles expressing all the tetraspanins simultaneously and with relatively high expression levels. Since they clustered together with the vesicles only expressing CD9-CD63, the results suggested a possible different biological function of this subgroup of vesicles and/or a different biogenesis route, e.g. from plasma or endosome membrane. This behavior has also been hypothesized by other studies on EV markers.^[36–38]^

Moreover, the results also seem to reveal that the analyzed tetraspanins do not follow random distributions but might be encapsulated into the vesicles according to different mechanisms. This behavior can be deduced from Figure 3F. A random distribution would result in the population of sEVs expressing just one of the markers to be the dominant one. This would be followed by those sEVs carrying two of the markers and lastly, by the population containing all three markers simultaneously. However, the data do not follow this trend for the HEK293 sEVs. In particular, 43% of the EV population in HEK293 cell line showed one marker only. This was followed by the EV population expressing three markers simultaneously (40%) and EVs expressing only two markers (17%). For the cbMSC and THP1 sEVs instead, most of EVs (>70%) expressed one marker only and were followed by EVs expressing two (~20%) and three markers (<10%). The same conclusion can be derived from the results of the combined FL-AFM measurements. A random tetraspanin distribution would also be likely accompanied by an increase in protein expression with sEV size, because of the larger surface area. Although the study by Tian et al. suggested this trend on EVs derived from colorectal cancer cells,^[13]^ our results showed no linear correlation between the FL intensity and the vesicle equivalent diameters. This behavior has also been observed in other investigations,^[37]^ indicating the possibility for selective EV cargo loading and/or different routes of biogenesis.^[37]^ Furthermore, a different correlation between the size and protein expression level may also be the result of differences in the EV isolation procedures and/or cellular source.

The AFM data of the vesicle sizes, combined with the FL data, also reveal that although not all the detected particles below 50 nm are EVs (these sizes are also detected in control substrates), there exist a population in this size range (Figure S10) that can be either very small EVs or exomeres.^[34]^ The comparison of the AFM results on the vesicle size and YM also suggested the presence of some common and distinct features among the samples. For example, the sub-population of HEK293 and cbMSC derived EVs that contains all the three tetraspanins simultaneously are mostly distributed around 80-90 nm (Figure S15). Furthermore, it seems that the high abundance of tetraspanins softens the average vesicle YM for these samples (Figure 5B cluster 1 for HEK293 sEVs and clusters 1 and 2 for cbMSC sEVs). The different behavior of the THP1 sEVs might be explained by both their lower abundance of vesicles/vesicle expressing tetraspanins (compared to the other cell lines) and by their low number of vesicles co-expressing the three tetraspanins simultaneously. Finally, the YM results seem to suggest that the stiffness of the analyzed vesicles is somehow affected by the abundance of the membrane tetraspanins, as also mentioned in other studies,^[21]^ even though their exact roles are not fully known.

## 4. Conclusion

In conclusion, we present a platform for high resolution and multiparametric analysis of single sEVs to resolve their heterogeneity. The platform combines atomic force and fluorescence microscopy to characterize individual sEVs and profile them based on their membrane protein composition, size and Young Modulus. The method applied to sEVs collected from the three different cell lines HEK293, cbMSC and THP1, shows the existence of several subpopulations bearing distinct properties, essentially highlighting the high EV heterogeneity. The data shows both common and cell-line specific features in the sEV populations. Although the sEVs investigated here are distributed in a large diameter range of 30-200 nm, the results do not show any size dependence of the expression levels of the common tetraspanins (CD9, CD63, CD81). However, some of the cell lines show a subpopulation of sEVs that simultaneously expresses all the tested tetraspanins in a relatively high abundance but existing mostly in a narrow size range of 70-100 nm. The level of CD9 and CD63 expressions in sEVs was also found to be strongly related to each other, irrespective of their cellular origin. On the contrary, the relative population of sEVs containing different combinations of the tested tetraspanins were found to depend on the cell line they are derived from. The values of Young Modulus also revealed some interesting features that could be related to their membrane protein composition and sizes. Although the biological significance of these observations is yet to be determined, the combined method shows a significant step forward for analysis of single vesicles. This combination of properties may provide new insights when analyzing EVs from clinical samples, potentially allowing to find new/more accurate diagnostic and/or therapeutic agent as well as help in deciphering unsolved questions regarding EV biology.

## 5. Experimental Methods

### Reagents

High purity deionized water (DIW) with a resistivity of 18 MΩ·cm was used throughout all the experiments. Phosphate-buffered saline (PBS, P4417) in tablets was purchased from Sigma-Aldrich. Anti-CD81-APC (A87789) was purchased from Beckman Coulter; anti-CD63-R-PE (1P-343-T100) and anti-CD9-PacificBlue (PB-208-T100) were purchased from ExBio. If not stated otherwise, all of the other chemicals were purchased from Sigma-Aldrich.

### EV purification and isolation

sEVs were prepared from THP-1 human monocytic cells, HEK293 Freestyle suspension cells (ThermoFisher), and immortalized human cord blood-derived mesenchymal stromal cells (cbMSCs; ATCC PCS-500-010) through ultrafiltration and bind-elute size exclusion chromatography (BE-SEC) as described previously.^[39]^ In brief, conditioned media were pre-cleared by low-speed centrifugation (5 min at 700 x g, then 10 min at 2000 x g) and by filtration through 0.22 **μ**m filters (Corning, cellulose acetate, low protein binding) before they were concentrated and diafiltrated with two times the initial volume of PBS by tangential flow filtration (300 kDa MidiKros columns, 370 cm^2^ surface area, Spectrum Labs). sEVs were further concentrated by using Amicon Ultra-15 10 kDa molecular weight cut-off spin filters (Millipore). Particle concentrations were assessed by NTA with a NanoSight NS500 instrument, as previously described.^[27]^ Fluorescently labelled sEVs were prepared accordingly from HEK293 Freestyle cells which were lentivirally engineered to stably express CD63-mNeonGreen fusion proteins as described before.^[27]^

### EV bulk surface characterization

The general surface marker composition for the sEVs from mNG-CD63 HEK293 cells was assessed by Western Blot (WB). The results showed the presence of common exosomal markers, e.g. CD9, CD63 and CD81, and the absence of other markers, e.g. GM130 marker typical for apoptotic bodies (data not shown). The general surface marker composition for the sEVs from the three cell sources (HEK293, cbMSC, THP1) was assessed by a multiplex bead-based flow cytometry assay (MACSPlex Exosome Kit, human, Miltenyi Biotec) by using an optimized workflow as described previously.^[27]^ In brief, 1 × 10^9^ NTA-based particles from each sample were incubated overnight in 96-well filter plates (supplied with the kit) with MACSPlex Exosome Capture Beads (5 *μ*L) on an orbital shaker at 450 RPM at room temperature. Beads were washed with MACSPlex buffer (200 *μ*L) and counterstained with a 1:1:1 mixture of APC-conjugated anti-CD9, anti-CD63 and or anti-CD81 Pan Tetraspanin detection antibodies (supplied in the kit, 4 *μ*L each) in a total volume of 135 *μ*L. The plate was incubated at 450 RPM for 1 h at room temperature. Next, the samples were washed twice in PBS and resuspended in MACSPlex buffer (150 *μ*L). Samples were then transferred to a V-bottom 96-well microtiter plate (Thermo Scientific) and analyzed by flow cytometry using a MACSQuant Analyzer 10 flow cytometer (Miltenyi Biotec). FlowJo software (version 10.6.2, FlowJo, LLC) was used to analyze flow cytometric data. Median fluorescence intensities (MFI) for capture bead subsets were background-corrected by subtracting respective MFI values from matched non-EV containing buffer controls that were treated exactly like EV samples (buffer + capture beads + antibodies). All incubation steps were performed in the dark. The characterization results are presented in Figure S2.

### FL analysis setup

The fluorescence (FL) measurements on the engineered mNG-EVs were performed with the 100x oil immersion lens of a Zeiss inverted microscope Axio Observer Z1, equipped with a thermoelectrically cooled CCD camera (Andor iXon3 888 EMCCD, −100 °C). The microscope included a 455 nm centered LED and a GFP-1828A-000 filter set (Semrock) for excitation of the mNG-CD63 protein. The images were acquired with the following settings: 1 MHz at 16 bit, EMG20.

The FL measurements on the wt-EVs were performed with the 100x oil immersion lens of a Zeiss inverted microscope Colibri 5, equipped with a Hamamatsu CCD Camera (Orca Flash 4). The microscope was equipped with four LEDs, centered at 385 nm, 475 nm, 555 nm, and 630 nm, respectively. All LEDs included individual excitation filters for correct fluorophore excitation and minimal crosstalk (Figure S1). The images were acquired with the following settings: 60% Fieldstop and 30% Aperture. The 475 nm, 555 nm and 630 nm LEDs were set at 100% power, whereas the 385 nm LED was set at 60% power to reduce image background and noise.

All the images, for both engineered and wild type EVs, were acquired using 2s acquisition time. All the results were collected for the EVs included in a microscope area (133.12 *μ*m ×133.12 *μ*m for 100x objective lenses).

### AFM setup

The AFM measurements were performed in liquid conditions on a JPK AFM system (NanoWizard® 3), under Quantitative Imaging (QI) Mode. QI is a force spectroscopy-based imaging mode that records a complete force distance curve at each pixel of the image, providing height information, but also adhesion and mechanical properties of the sample, i.e. Young Modulus. As there are nearly no lateral forces involved, this imaging mode is particularly suitable for soft samples, such as cells and single particles, i.e. EVs. For the measurements, quartz-like NANOSENSORS qp-BioAC-CI AFM probes (CB2 cantilever, 0.1 Nm^−1^ force constant) were used.

The scanning parameters, including Setpoint, Z-length, Pixel Time, Z-range and N° of pixels, were optimized for different images, depending on their desired resolution, and were kept constant for the same image types. The threshold height used for EV identification was 15 nm.

### Single EV analysis platform

Experiments were performed on a combined inverted microscope and AFM setup aligned with the optical axis of the microscope, using 170 *μ*m thick coverslips from Ibidi (Gridded Glass Coverslips Grid-50). The coverslips included four imprinted grids for area identification and were further patterned with marks (equidistant groups of four crosses per group, 5*μ*m long and 50-60 nm deep), using focused ion beam (FIB). This step was necessary for precise area identification and overlap of the FL and AFM images.

sEVs were first covalently captured onto a coverslip inserted in a chamber well, following our previously reported functionalization protocol.^[26]^ Briefly, the coverslip was first cleaned in a 5:1:1 solution of DI water, H_2_O_2_ and NH_4_OH (88°C, 10 min) and activated with (3-aminopropyl)triethoxysilane (APTES, 5% v/v in 95% ethanol, 10 min) and glutaraldehyde (GA, 1% v/v in 1x PBS, 1 h). Thereafter, EVs were covalently immobilized on the GA surface for 1 h. For the engineered mNG-EV measurements, this step was followed by PBS washing, which completed the functionalization. For the wild type EVs instead, the EV capture step was followed by deactivation of the remaining active groups with Tris-Ethanolamine (Tris-ETHA, 0.1 M Tris buffer and 50 mM ethanolamine, pH 9.0, 30 min) and casein (0.05% w/v in 1x PBS, 90 min) in order to minimize non-specific binding (NSB) of the fluorescently labelled antibodies. Finally, the coverslips with immobilized sEVs were incubated with a solution containing the three fluorescently labelled CD81-APC, CD63-R-PE and CD9-PB antibodies (2 nM each in 1x PBS, for 2 h) and washed with 1xPBS for the subsequent analyses. Being the EVs immobilized at a concentration of 0.33 pM, they were incubated with an antibody concentration more than 6000-fold in excess to guarantee proper tetraspanin staining. All the FL and AFM measurements were performed in liquid, in 1x PBS.

Following functionalization, the chamber well was positioned on top of the inverted microscope and the AFM tip on top was then aligned with the microscope optical axis, in order to scan the same fluorescently imaged area. After alignment, FL images with the different LEDs were taken, followed by AFM scanning. The presence of the FIB marks on the coverslips was essential in this phase, as it could overcome for small shifts between the FL and AFM scans, occurring although the initial tip-lens alignment step, and enabled the accuracy of the combined measurements.

The AFM scans were first taken on top of the FIB marks for big spots (30 *μ*m × 30 *μ*m) at low resolutions, for area identification, and then for small spots (5 *μ*m × 5 *μ*m) at high resolution (256 × 256 pixels, 19.5 nm pixel resolution), for accurate EV size determination.

### FL image processing and analysis

The FL images were processed and analyzed using the Zeiss Image Analysis Software (ZEN, Blue Edition). Briefly, after background generation and subtraction, the “Image overlay” processing function was used to stack the FL images of the different channels (different antibodies) together. Thereafter, a custom analysis workflow (Analysis window – segment region classes independently method) containing appropriate values of intensity threshold, smoothing, dot separation and area for each channel was created and applied to the stacked image. For each fluorescent spot, different parameters including the integrated intensity, pixel count, etc. were extracted and analyzed. The same workflow was applied for the three different cell-line derived sEVs.

A custom-made MATLAB script was utilized to consolidate the data files generated by the ZEN software for each fluorescence channel, to calculate the total numbers of EVs detected and classify the EVs based on the coexistence of various markers.

### AFM image processing and analysis

The AFM height images were processed using the JPK data processing software. First, a custom workflow containing plane/line fit corrections and a median filter was applied to all images. This allowed the removal of noise and artifacts coming from substrate tilting. Following correction, Gwyddeon software and a MATLAB code were used for analysis. In particular, a maximum height threshold of 15 nm was set for EV identification and the surface area of all the particles above threshold was extracted and analyzed. Assuming surface area conservation, the EV equivalent diameters were then obtained for single vesicles.

The AFM Young Modulus images were processed using the JPK data processing software. In particular, a custom workflow containing baseline subtraction, contact point determination, vertical tip position calculation and elasticity fit was applied to all the force curve files of the scanned areas. Following correction, the Young Modulus image data were analyzed using a custom MATLAB code. In particular, a mask based on each AFM Height image, where the EVs were identified, was generated using Gwyddeon. The mask was then transferred to the corresponding AFM Young Modulus image and the mean Young Modulus for each vesicle was extracted. Finally, a script in MATLAB was used to connect each EV to its mean Young Modulus value.

### tSNE analysis

For an easier overlook of the acquired data, the FL integrated intensities of the three analyzed tetraspanins of the EVs were mapped onto a two-dimensional plane using tSNE. Different tSNE mappings with perplexity values ranging from 1-40 were tested. The optimal value of perplexity was determined by investigating each perplexities’ pseudo-BIC value.^[40]^ The maximum number of iterations in the algorithm was kept sufficiently large to ensure convergence. As the parameters were continuous, the tSNE algorithm was run with standardized Euclidean distance metric. After mapping the data with tSNE, cluster algorithms were applied on the result, making it possible to reveal subpopulations of the vesicles which could be analyzed. The number of clusters was determined by visual inspection of the tSNE plots for each cell-line. Each vesicle was then sorted into one of the identified clusters by applying a linkage hierarchical clustering algorithm. For HEK293 a tSNE mapping with a perplexity of 29 and with 5 identified clusters was selected, and for cbMSC a tSNE mapping with a perplexity of 32 and with 3 identified clusters was selected. For the third cell line, THP1, a tSNE mapping with a perplexity of 17 and with 5 identified clusters was selected. Both the tSNE and the cluster analysis were performed using a MATLAB script.

## Supporting Information

Supporting Information is available from the website or from the author.

## Acknowledgements

The entire study was supported by a grant from the Erling Persson Family Foundation. A.D., F.P. and S.S.S. acknowledge also the grant funded by the Swedish Research Council (Contract No. 2016-05051). SELA is supported by the Swedish Research Council and SSF-IRC (FormulaEx). The authors acknowledge the work of Lars Riekehr (Uppsala University, Sweden), who patterned the marks needed for precise spot identification on the coverslips used for combined FL-AFM measurements, using FIB. They also acknowledge Marco Colangelo (MIT, Boston) for the help in the MATLAB coding. Figure 1 was realized using BioRender.com. AG is an International Society for Advancement of Cytometry (ISAC) Marylou Ingram Scholar 2019-2023). DRM acknowledges the support from Cihan University-Erbil, in Iraq. Authors contributed as follows. SC conceptualized the study, performed the combined measurements on the mNG-EVs (together with CP) and the measurements on wt-EVs, performed the control measurements, processed, analyzed and plotted the data, wrote the original manuscript and designed the figures under the supervision of AD. FP conceptualized the study, developed the combined FL-AFM platform, performed the measurements on the stability of the vesicles and on the capture efficiency of the protocol. FS extracted the size and YM information from the AFM data, performed the tSNE and cluster analysis on the wt-EVs. AG advised on study design, coordinated production, isolation and characterization of EVs, performed the multiplex bead-based flow cytometry EV surface profiling experiments and analyzed the corresponding data. CP performed the FL-AFM combined measurements on the mNG-EVs under the supervision of FP and SC. SSS wrote the MATLAB script to consolidate the FL data and classify EVs based on the co-existence of various markers, contributed to the tSNE analysis. DRM contributed to isolate and purify the sEV samples and to characterize them by NTA. DG contributed to characterize the sEV samples by NTA and WB. SELA advised on study design and led the lab generating all EVs for the study. JL reviewed and edited the manuscript, acquired the funding. AD conceptualized the study, analyzed the data, supervised the study, wrote the original manuscript. All authors reviewed and have given approval to the final version of the manuscript.

## TOC

Extracellular vesicles (EVs) offer unique opportunities for both diagnostics and therapeutics. In this paper, the advantages of using a combined fluorescence and atomic force microscopy for multiparametric profiling of single small EVs is presented. By studying and correlating membrane protein compositions, size and mechanical properties of single small EVs, the platform can provide new insights about EV subpopulations and heterogeneity.

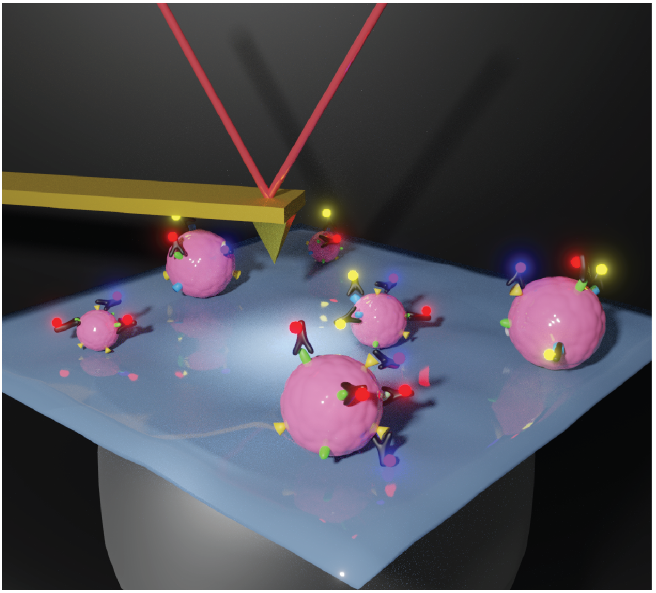

## References

[1] D. E. Murphy, O. G. De Jong, M. Brouwer, M. J. Wood, G. Lavieu, R. M. Schiffelers, P. Vader, Exp. Mol. Med. 2019, 51, 1.

[2] A. Louise, S. Revenfeld, R. Bæk, M. H. Nielsen, A. Stensballe, K. Varming, M. Jørgensen, Clin. Ther. 2014, 36, 830.

[3] P. Vader, X. O. Breakefield, M. J. A. Wood, Trends Mol. Med. 2014, 20, 385.

[4] S. El Andaloussi, I. Mäger, X. O. Breakefield, M. J. A. Wood, Nat. Rev. Drug Discov. 2013, 12, 347.

[5] E. Willms, H. J. Johansson, I. Mäger, Y. Lee, K. E. M. Blomberg, M. Sadik, A. Alaarg, C. I. E. Smith, J. Lehtiö, S. E. L. Andaloussi, M. J. A. Wood, P. Vader, Sci. Rep. 2016, 6, 1.

[6] L. M. Id, Y. Sadovsky, PLoS Biol. 2019, 17, 1.

[7] C. Lässer, S. Chul, L. Jan, Mol. Aspects Med. 2018, 60, 1.

[8] D. K. Jeppesen, A. M. Fenix, J. L. Franklin, J. N. Higginbotham, Q. Zhang, L. J. Zimmerman, D. C. Liebler, J. Ping, Q. Liu, R. Evans, W. H. Fissell, J. G. Patton, L. H. Rome, D. T. Burnette, R. J. Coffey, Cell 2019, 177, 428.

[9] T. L. Whiteside, Expert Rev. Mol. Diagn. 2015, 15, 1293.

[10] S. L. N. Maas, X. O. Breakefield, A. M. Weaver, M. G. Hospital, Trends Cell Biol. 2017, 27, 172.

[11] T. A. Hartjes, S. Mytnyk, G. W. Jenster, V. Van Steijn, Bioengineering 2019, 6, 7.

[12] K. Lee, K. Fraser, B. Ghaddar, K. Yang, E. Kim, L. Balaj, E. A. Chiocca, X. O. Breake, H. Lee, R. Weissleder, ACS Nano 2018, 12, 494.

[13] Y. Tian, M. Gong, G. Su, S. Zhu, W. Zhang, S. Wang, Z. Li, C. Chen, L. Li, L. Wu, X. Yan, ACS Nano 2018, 12, 671.

[14] G. G. Daaboul, P. Gagni, L. Benussi, P. Bettotti, M. Ciani, M. Cretich, D. S. Freedman, R. Ghidoni, A. Y. Ozkumur, C. Piotto, D. Prosperi, B. Santini, M. S. Ünlü, M. Chiari, Sci. Rep. 2016, 6, 1.

[15] R. P. Carney, S. Hazari, M. Colquhoun, D. Tran, B. Hwang, M. S. Mulligan, J. D. Bryers, E. Girda, G. S. Leiserowitz, Z. J. Smith, K. S. Lam, Anal. Chem. 2017, 89, 5357.

[16] D. Wu, J. Yan, X. Shen, Y. Sun, M. Thulin, Y. Cai, L. Wik, Q. Shen, J. Oelrich, X. Qian, K. L. Dubois, K. G. Ronquist, M. Nilsson, U. Landegren, M. Kamali-moghaddam, Nat. Commun. 2019, 10, 1.

[17] G. Corso, W. Heusermann, D. Trojer, A. Görgens, E. Steib, J. Voshol, A. Graff, C. Genoud, J. Hean, J. Z. Nordin, O. P. B. Wiklander, S. El Andaloussi, N. Meisner-kober, G. Corso, W. Heusermann, D. Trojer, A. Görgens, J. Voshol, A. Graff, C. Genoud, Y. Lee, J. Hean, Z. Joel, O. P. B. Wiklander, S. El Andaloussi, N. M. Systematic, J. Extracell. Vesicles 2019, 8, 1663043.

[18] B. . Görgens, A.; Bremer, M.; Ferrer-Tur, R.; Murke, F.; Tertel, T.; Horn, P.A.; Thalmann, S.; Probst, C.; Guerin, C.; Boulanger, C.M.; Hanenberg, H.; Erdbruegger, U.; Lannigan, J.; Ricklefs, F.L.;Andaloussi, S. El; and Giebel, J Extracell Vesicles, Press. 2019, 8, 1587567.

[19] F. A. W. Coumans, A. R. Brisson, E. I. Buzas, F. Dignat-george, E. E. E. Drees, S. El-andaloussi, C. Emanueli, A. Gasecka, A. Hendrix, A. F. Hill, R. Lacroix, Y. Lee, T. G. Van Leeuwen, N. Mackman, I. Mäger, J. P. Nolan, E. Van Der Pol, D. M. Pegtel, S. Sahoo, P. R. M. Siljander, G. Sturk, O. De Wever, R. Nieuwland, Circ. Res. 2017, 120, 1632.

[20] M. Yáñez-Mó, P. R.-M. Siljander, Z. Andreu, A. Bedina Zavec, F. et al Borràs, J. Extracell. Vesicles 2015, 4, 27066.

[21] D. Vorselen, S. M. Van Dommelen, R. Sorkin, M. C. Piontek, J. Schiller, S. T. Döpp, S. A. A. Kooijmans, B. A. Van Oirschot, B. A. Versluijs, M. B. Bierings, R. Van Wijk, R. M. Schiffelers, G. J. L. Wuite, W. H. Roos, Nat. Commun. 2018, 9, 1.

[22] R. Sorkin, R. Huisjes, F. Boškovic, D. Vorselen, S. Pignatelli, Y. Ofir-birin, J. K. F. Leal, J. Schiller, D. Mullick, W. H. Roos, G. Bosman, N. Regev-rudzki, R. M. Schiffelers, G. J. L. Wuite, Small 2018, 14, 1.

[23] L. Zhang, Q. Feng, J. Wang, S. Zhang, B. Ding, Y. Wei, M. Dong, J. Ryu, T. Yoon, X. Shi, J. Sun, X. Jiang, ACS Nano 2015, 9, 9912.

[24] E. Van der Pol, F. A. Coumans, A. Grootemaat, C. Gardiner, I. Sargent, P. Harrison, A. Sturk, T. Van Leeuwen, R. Nieuwland, J. Thromb. Haemost. 2014, 12, 1182.

[25] S. Sharma, M. Leclaire, J. K. Gimzewski, Nanotechnology 2018, 29, 132001.

[26] S. Cavallaro, J. Horak, P. Hååg, D. Gupta, C. Stiller, S. S. Sahu, A. Görgens, H. K. Gatty, K. Viktorsson, S. El Andaloussi, R. Lewensohn, A. E. Karlstrom, J. Linnros, A. Dev, ACS Sensors 2019, 4, 1399.

[27] A. Wiklander, O.P.B; Bostancioglu, R.B.; Welsh, J.A.; Zickler, A.M.; Murke, F.; Corso, G.; Felldin, U.; Hagey, D.W.; Evertsson, B.; Liang, X.M.; Gustafsson, M.O.; Mohammad, D.K.; Wiek, C.; Hanenberg, H.; Bremer, M.; Gupta, D.; Björnstedt, M.; Giebel, B.; No, Front. Immunol. 2018, 9.

[28] M. Reth, Nat. Immunol. 2013, 14, 765.

[29] J. Syed, J. Ashton, J. Joseph, G. N. Jones, C. Slater, A. Sharpe, G. Ashton, W. Howat, R. Byers, H. K. Angell, Immunother. Open Access 2019, 5, DOI 10.35248/2471-9552.19.5.157.

[30] G. Corso, I. Mäger, Y. Lee, A. Görgens, J. Bultema, B. Giebel, M. J. A. Wood, J. Z. Nordin, S. E. L. Andaloussi, Sci. Rep. 2017, 7, 11561.

[31] D. Bachurski, M. Schuldner, P. Nguyen, A. Malz, K. S. Reiners, J. Extracell. Vesicles 2019, 8, DOI 10.1080/20013078.2019.1596016.

[32] A. Ridolfi, M. Brucale, C. Montis, L. Caselli, L. Paolini, A. Borup, A. T. Boysen, F. Loria, M. J. C. Van Herwijnen, M. Kleinjan, P. Nejsum, N. Zarovni, M. H. M. Wauben, D. Berti, P. Bergese, F. Valle, Anal. Chem. 2020, 92, 10274.

[33] D. Kim, H. Nishida, S. Y. An, A. K. Shetty, T. J. Bartosh, D. J. Prockop, Proc. Natl. Acad. Sci. 2015, 113, 170.

[34] Q. Zhang, J. N. Higginbotham, D. K. Jeppesen, Y. Yang, W. Li, T. Mckinley, R. Graves-deal, J. Ping, C. M. Britain, K. A. Dorsett, C. L. Hartman, D. A. Ford, R. M. Allen, K. C. Vickers, Q. Liu, L. Jeffrey, S. L. Bellis, R. J. Coffey, Cell Rep. 2019, 27, 940.

[35] Y. Yang, G. Shen, H. Wang, H. Li, T. Zhang, N. Tao, X. Ding, H. Yu, 2018, 115, 10275.

[36] J. Kowal, G. Arras, M. Colombo, M. Jouve, P. J. Morath, B. Primdal-bengtson, D. Florent, D. Loew, M. Tkach, C. Théry, Proc. Natl. Acad. Sci. 2016, 113, E968.

[37] F. K. Fordjour, G. G. Daaboul, S. J. Gould, bioRxiv 2019.

[38] Y. Ji, D. Qi, L. Li, H. Su, X. Li, Y. Luo, B. Sun, F. Zhang, B. Lin, T. Liu, Y. Lu, Proc. Natl. Acad. Sci. 2019, 116, 5979.

[39] G. Corso, I. Mäger, Y. Lee, A. Görgens, J. Bulte, B. Giebel, M. J. A. Wood, J. Z. Nordin, S. E. L. Andaloussi, Sci. Rep. 2017, 1.

[40] Y. Cao, L. Wang, arXiv Prepr. arXiv1708.03229 2017, 1.

