## Supporting Information for "Multiparametric Profiling of Single Nanoscale Extracellular Vesicles by Combined Atomic Force and Fluorescence Microscopy: Correlation and Heterogeneity in Their Molecular and Biophysical Features"

**Figure S1.**

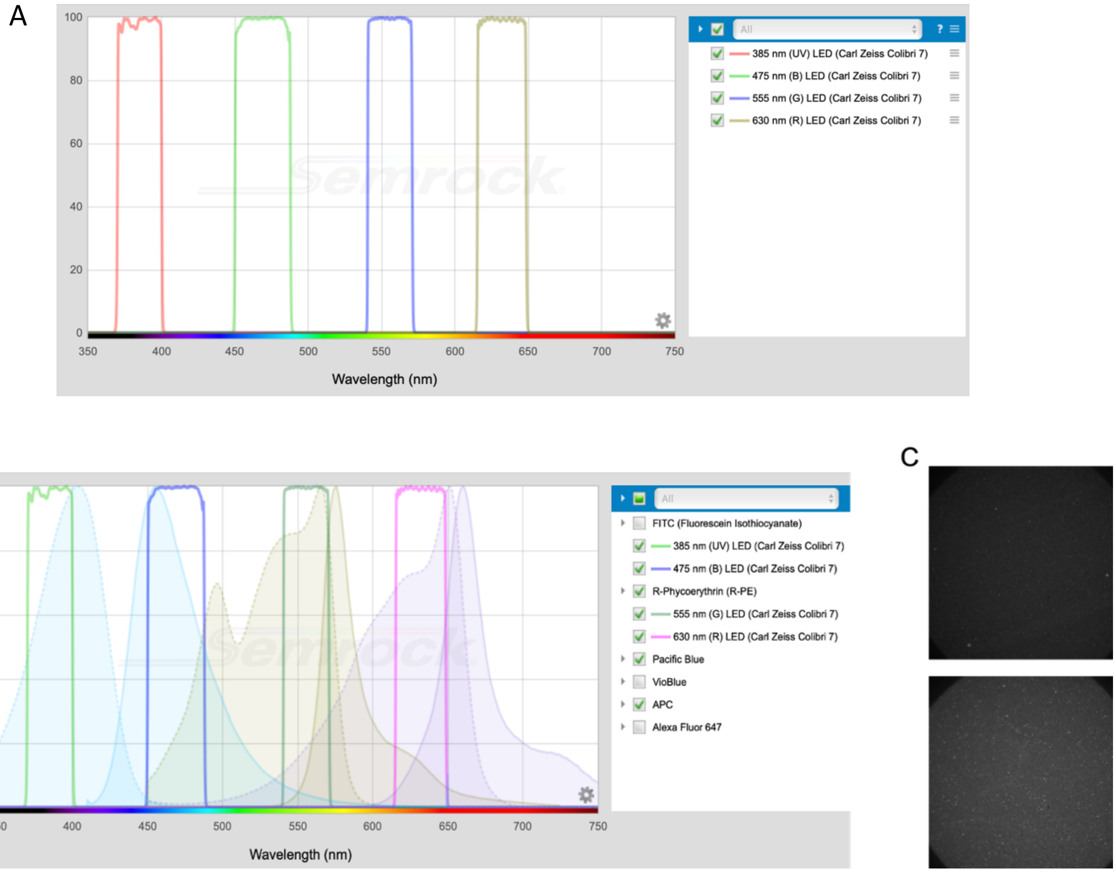


**Figure S1**. A) schematic of the LED sources and built-in excitation filters available in our inverted microscope, used to analyze the wt-EVs. B) Schematic of the excitation and emission spectra of the fluorescently labelled antibodies used to target the EV tetraspanins: CD81-APC, CD63-R-PE and CD9-PacificBlue. As shown, PacificBlue fluorophore is only excited by the 385nm LED. The R-PE fluorophore is only excited by the 555nm LED, while the APC fluorophore is excited by the 630 nm LED, with minimum cross-excitation with the 555 nm LED. (C) Comparison between the fluorescence spots generated by the CD81-APC antibody when excited with 555 nm LED (top) and the spots generated by CD63-R-PE fluorophore (bottom), when excited with the same source. As visible, the number of spots and intensity values are much lower for the CD81-APC antibody compared to the CD63-R-PE one, indicating negligible crosstalk in the 555 nm LED channel.

**Figure S2**.

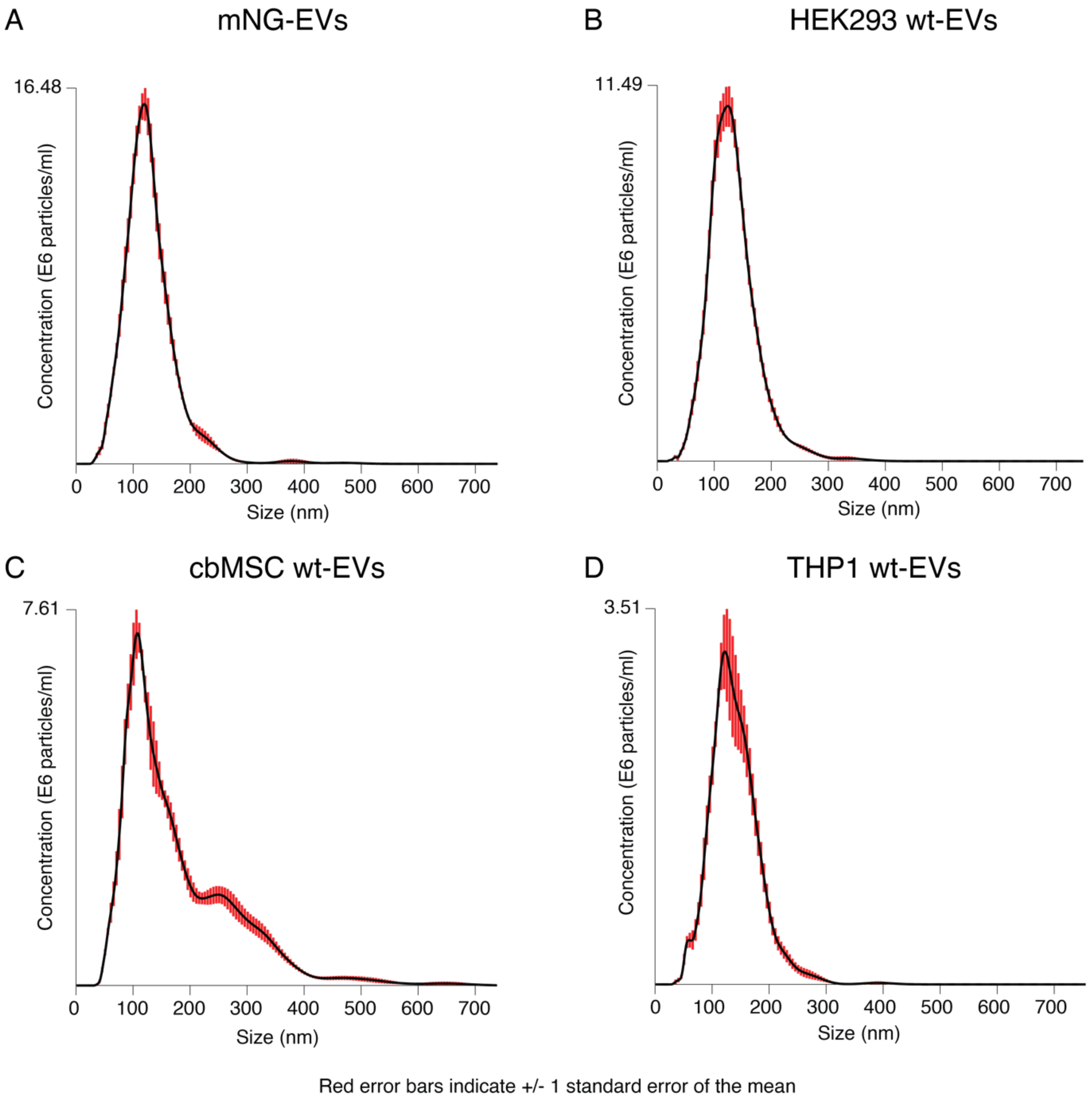

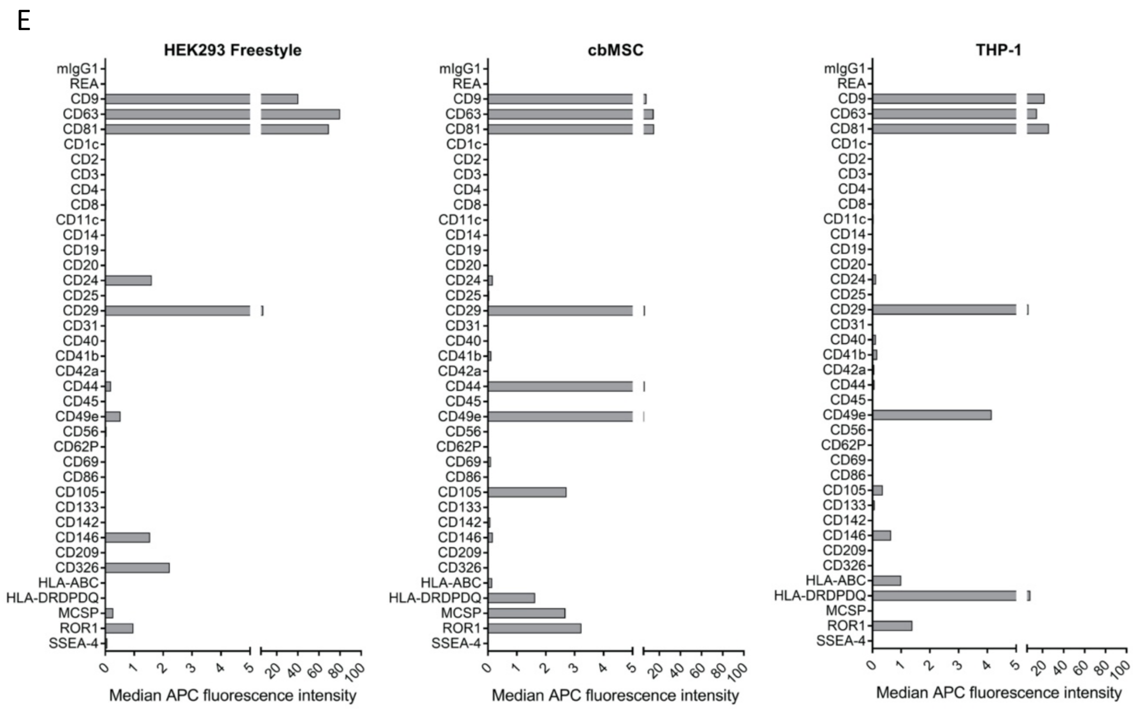


**Figure S2**. sEV characterization results. (A-D) NTA scatter-based size distribution profiles of the mNG-EVs from the HEK293 cell line, wt-EVs from HEK293, wt-EVs from cbMSC and wt-EVs from THP1 cell lines, respectively. (E) MACSPlex surface protein characterization of the three wt-EVs. As shown, all three tetraspanins CD9, CD63 and CD81 are expressed at a relatively high abundance on sEVs in all three samples. Other markers are expressed differentially between samples (e.g. CD105 on MSC-EVs, which is a known MSC marker).

**Figure S3**.

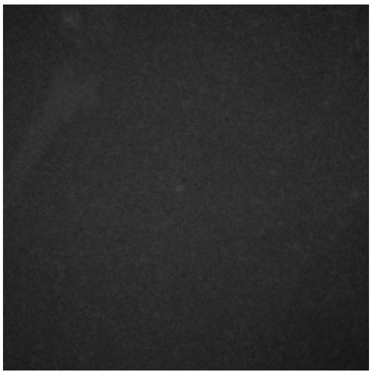

**Figure S3**. Control substrate functionalized up to glutaraldehyde (GA), without mNG-EVs. As shown, no bright FL spots were detected on the substrate.

**Figure S4**.

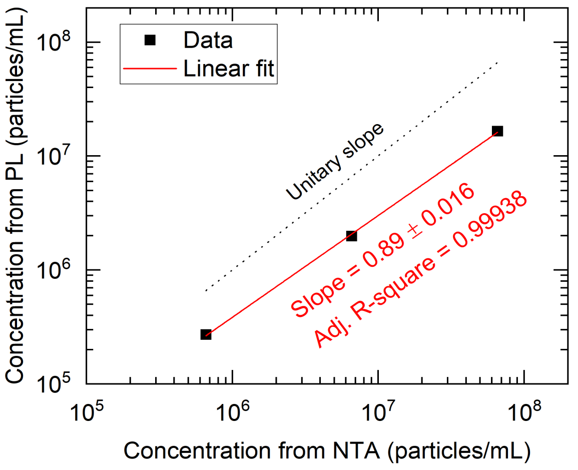


**Figure S4**. Correlation between the number of particles detected by NTA (x-axis) and those detected by our single EV platform (y-axis). Number of particles calculated considering homogeneous capture of the vesicles on the substrate.

**Figure S5**.

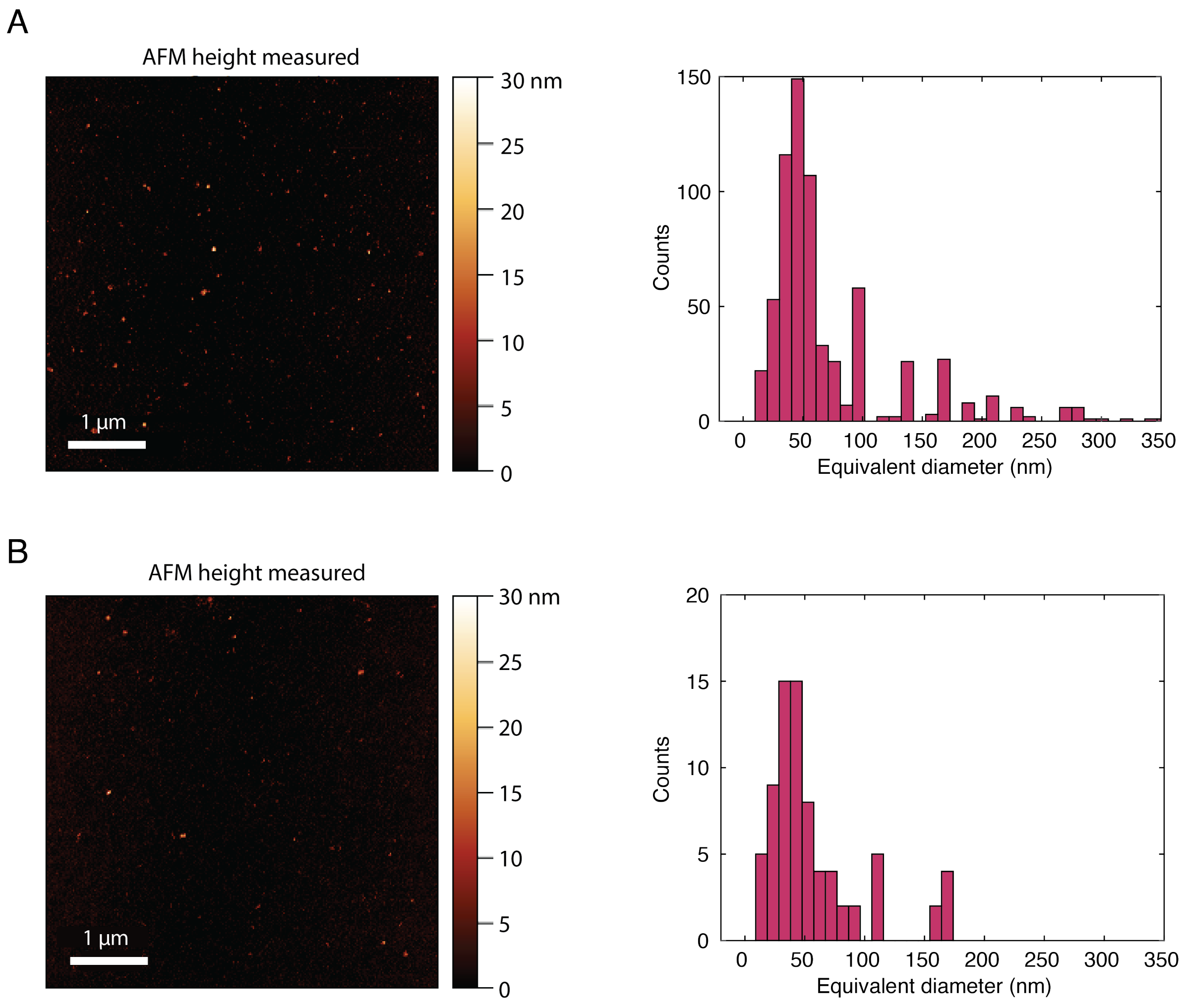

**Figure S5**. AFM scans of control surfaces. (A) Representative AFM scan of a control substrate functionalized up to GA, without sEVs, and corresponding equivalent diameter distribution of the control surfaces. As shown, the distribution is shifted towards smaller sizes compared to the surfaces with EVs, peaking at around 30-40 nm. Distribution obtained from 14 scans of 5 µm x 5 µm. (B) Representative AFM scan of a control surface functionalized up to GA and followed by storage buffer (SB) incubation (same dilution as buffer concentration in the sEV solution), tris-ETHA and casein. Scan followed by the corresponding equivalent diameter distribution of the control surfaces with SB. As before, the distribution is shifted towards small sizes, peaking at around 30-40 nm. Distribution obtained from 11 scans of 5 µm x 5 µm. Lower number of particles compared to the previous image because lower number of areas scanned, and the surface is more uniform due to the use of the blocking layer (tris-ETHA and casein).

**Figure S6**.

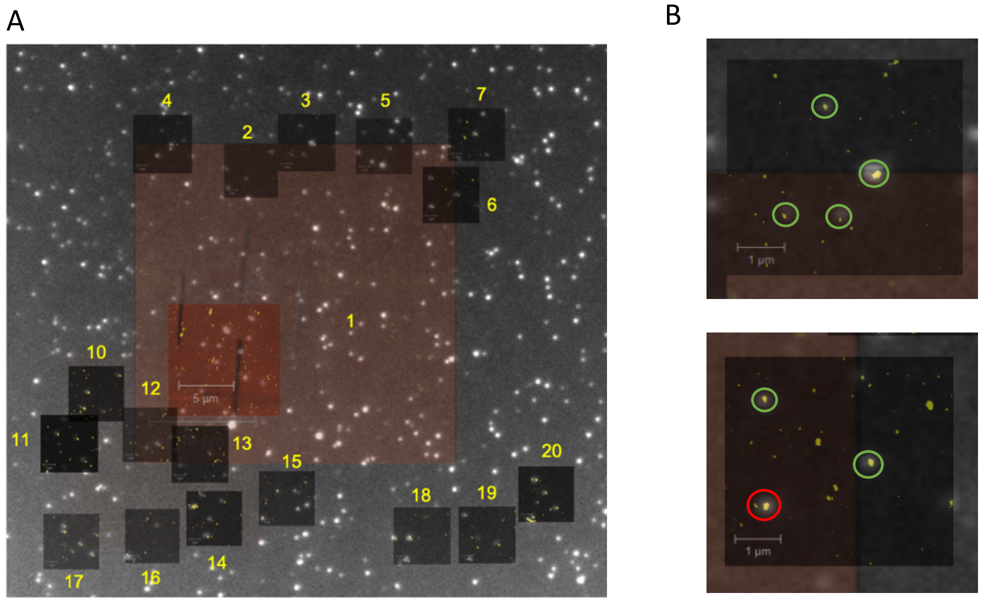


**Figure S6**. (A) Representative image of a combined FL-AFM measurement for mNG-EVs. The crosses patterned with FIB were visible in both FL and AFM scans, allowing precise overlapping of the two images and area/EV identifications. The bigger AFM scans (30 µm x 30 µm) were taken at a low resolution for marker identification, whereas the smaller AFM scans (5 µm x 5 µm) were taken at high resolution for precise EV size determination. (B) Zoom in of two areas with overlapping FL and AFM images. As shown, most of the bright diffuse FL spots (encircled in green) corresponded to single EVs (yellow particles in inner AFM scans), with only very few spots corresponding to multiple particles (encircled in red). These multiple particle spots were not considered for the single EV analysis.

**Figure S7**.

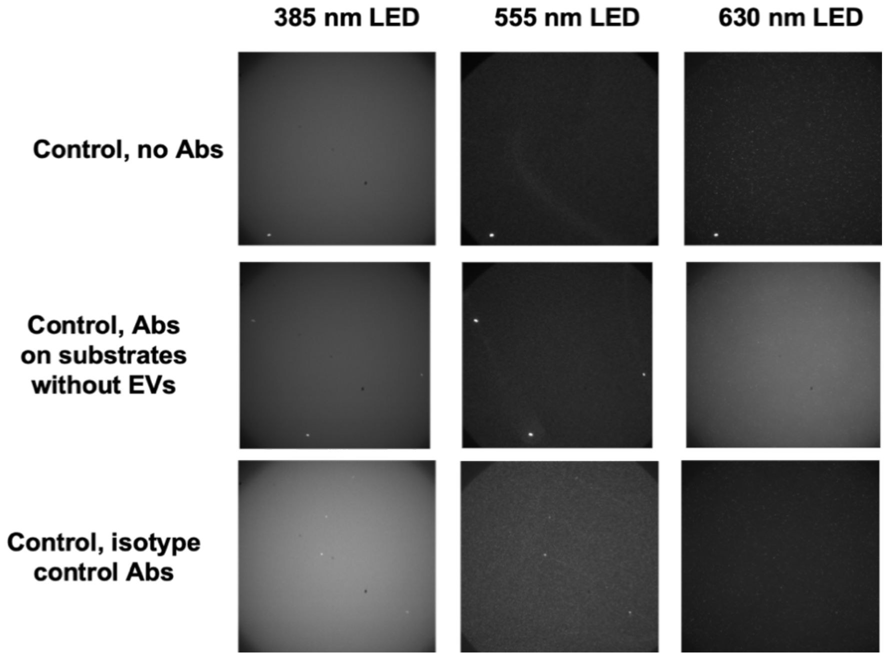


**Table 1**. Table showing the intensity values calculated over 6 different control substrate areas and expressed as CCD counts/px.

| Control substrate | 385 nm LED | 555 nm LED | 630 nm LED |
|---|---|---|---|
| Substrate with EVs, without antibodies | 5869 ± 88 | 659 ± 64 | 407 ± 44 |
| With CD81-APC, CD63-R-PE, CD9-PB Abs on substrate without EVs | 5934 ± 146 | 815 ± 72 | 1807 ± 97 |
| With isotype control Abs CD81-APC, CD63-R-PE, CD9-PB on substrate with EVs | 5888 ± 142 | 697 ± 81 | 495 ± 43 |

**Figure S7**. Control images acquired with the 385 nm, 555 nm and 630 nm LEDs corresponding to the PB, R-PE and APC fluorophores conjugated with the antibodies, respectively. First line: control substrate with EVs but without antibody staining. Second line: control substrate without EVs but with CD9-PB, CD63-R-PE and CD81-APC antibody staining. Third line: control substrate with isotype control antibody staining, where antibodies were labelled with the same PB, R-PE and APC fluorophores used for EV analysis. In all cases, the control substrates were functionalized following all the steps up to casein, including EV immobilization for the first and third cases. Table 1 reports the FL intensity values for all the three controls, showing that the values did not change significantly upon antibody incubation, therefore showing no unspecific antibody binding. Only the control with CD81-APC showed higher values than the others, indicating small unspecific binding. However, these values were still much lower than the EV intensity values and thus did not affect the measurements.

**Figure S8.**

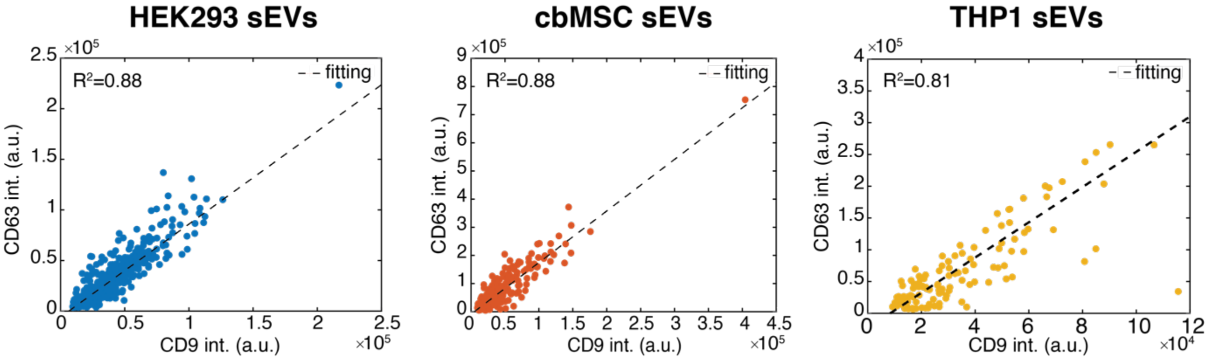


**Figure S8.** Plots showing the correlation of the CD9 and CD63 FL intensities for the CD9-CD63 only positive sEVs of HEK293, cbMSC and THP1 cell lines.

**Figure S9.**

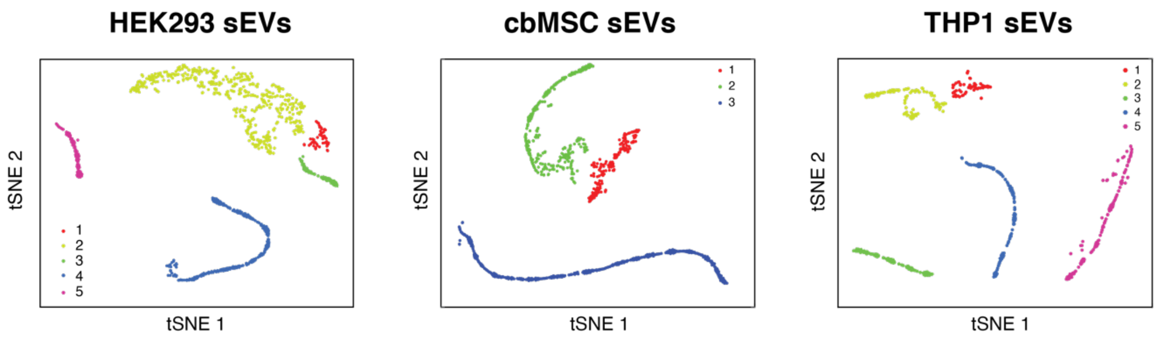


**Figure S9.** Two dimensional tSNE mapping of the sEV FL integrated intensity values showing 5, 3 and 5 optimized clusters for the HEK293, cbMSC and THP1 sEVs, respectively.

**Figure S10.**

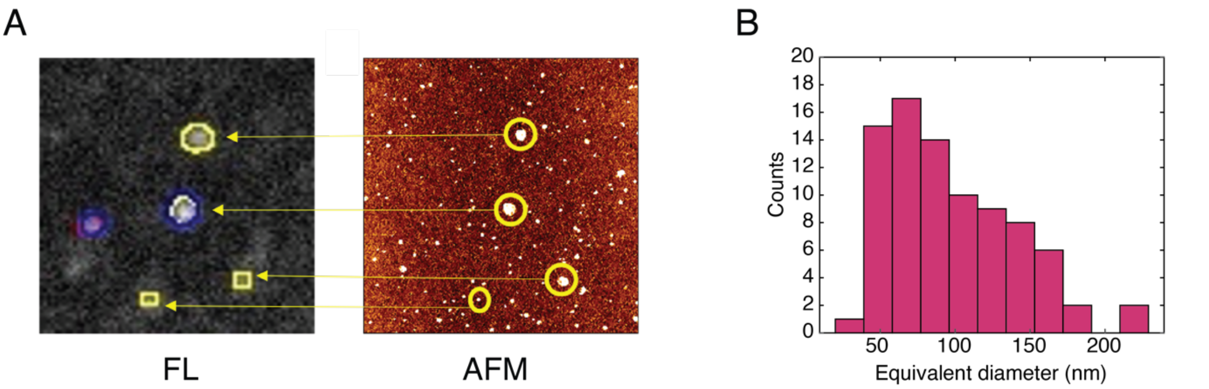


**Figure S10.** (A) Representative FL and AFM combined measurement performed on wt-EVs from HEK293 cell line. The AFM particles on the right image encircled in yellow are those corresponding to the FL particles on the left image expressing the different tetraspanins. (B)

Representative equivalent diameter distribution of the sEVs analyzed by AFM during the combined measurements (those for which FL was detected) for HEK293 cell line.

**Figure S11.**

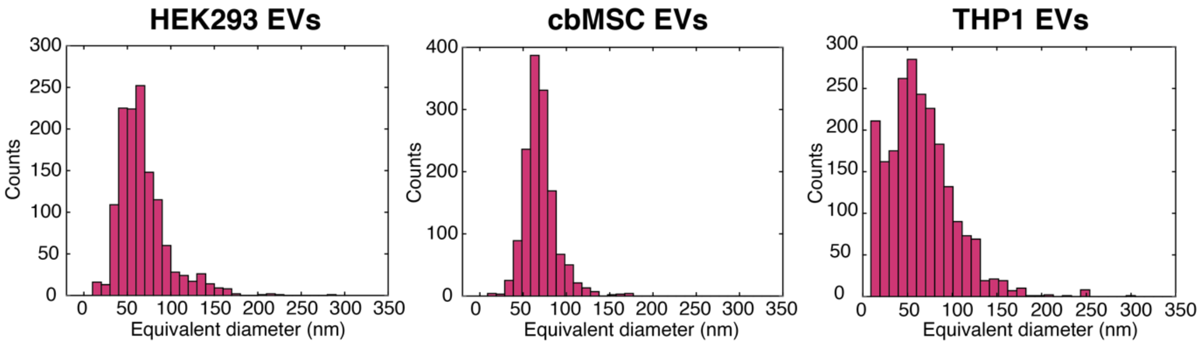


**Figure S11**. AFM equivalent diameter distributions for all the HEK293, cbMSC and THP1 sEVs analyzed by AFM during the combined measurements. The distributions also include the vesicles that did not show any FL signal (no tetraspanins or very low levels below the detection range).

**Figure S12**.

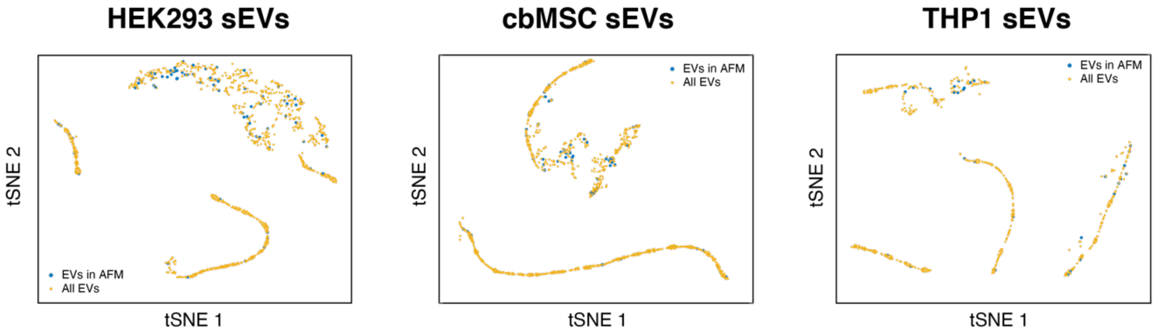


**Figure S12** Distribution of the sEVs analyzed by AFM (blue points) among all the clusters identified by the tSNE considering the sEVs analyzed by FL (yellow points). As shown, the vesicles are uniformly distributed among all the identified clusters.

**Figure S13.**

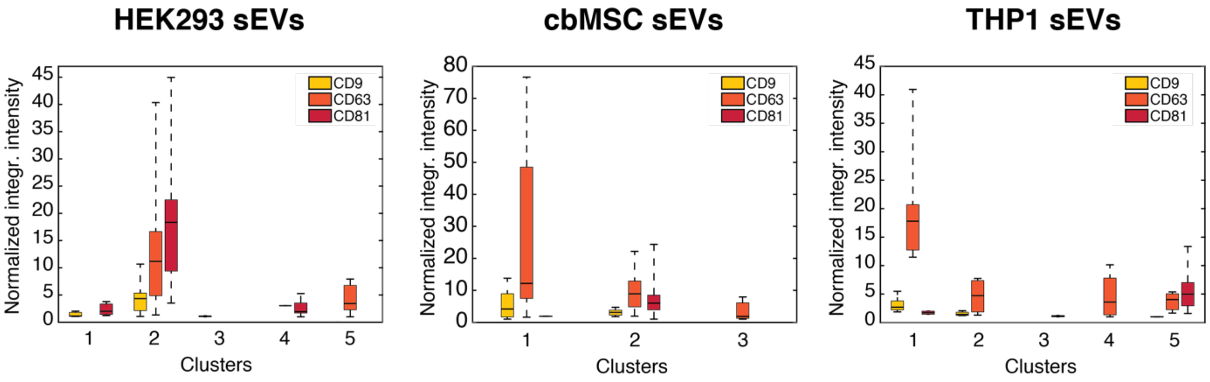


**Figure S13.** Box plots of the normalized FL intensity distributions of the three tetraspanins for the tSNE derived clusters for the sEVs analyzed by AFM. As shown, the tetraspanin intensity distribution is similar to that obtained for the whole sEV dataset reported in Figure 4B.

**Figure S14.**

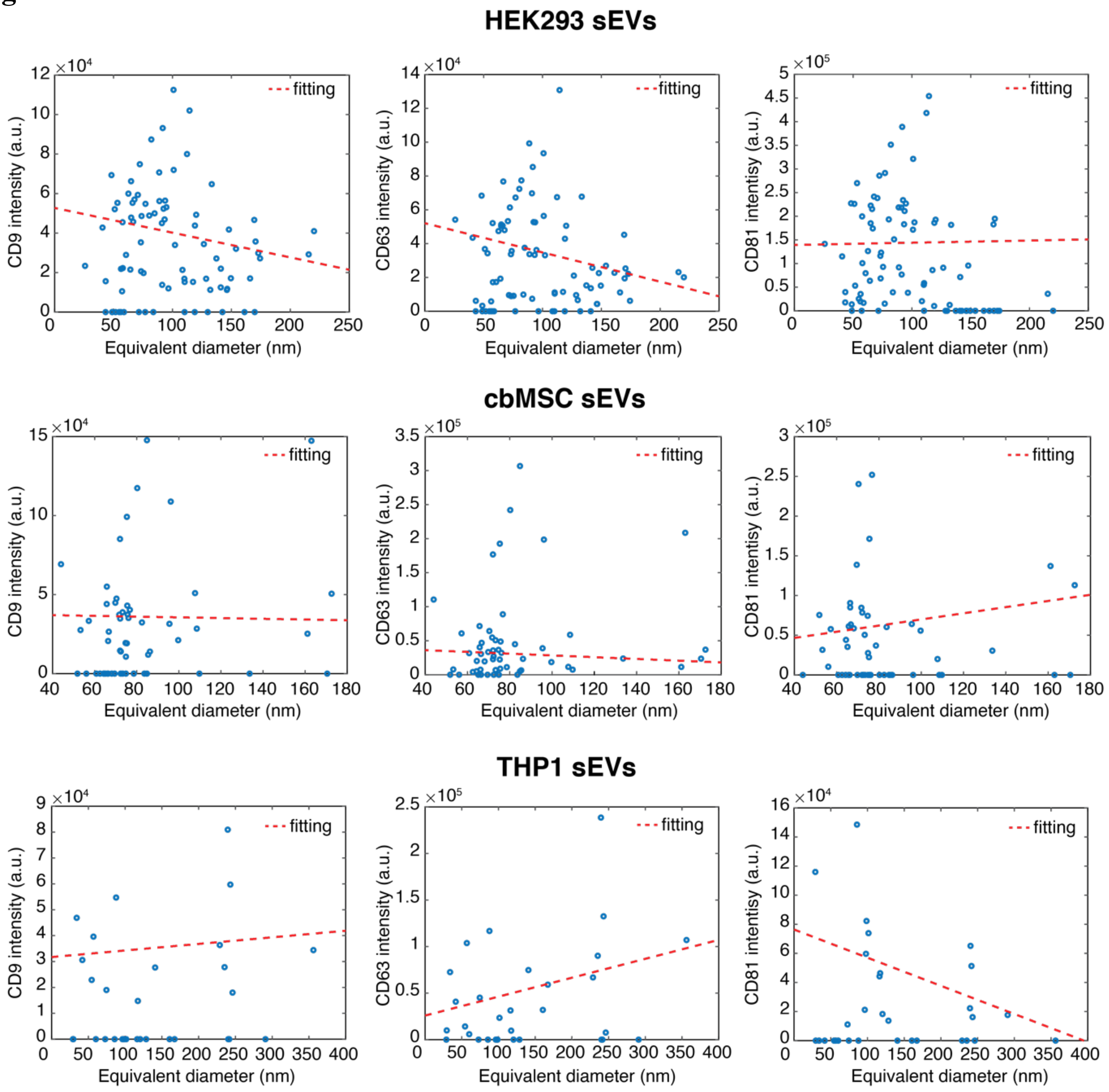

**Figure S14**. Correlation plots between the tetraspanin FL intensities and the sEV equivalent diameters for HEK293, cbMSC and THP1 EVs, respectively. As shown by the poor fitting, the protein expressions on the sEVs were not linearly correlated to their sizes, indicating that a higher amount of tetraspanins did not necessarily mean a bigger vesicle.

**Figure S15**.

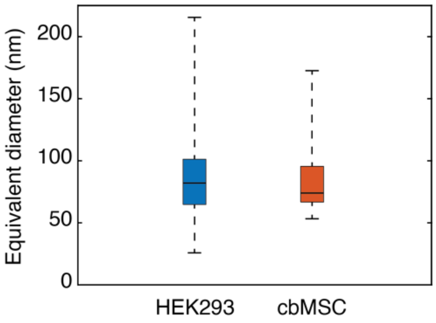


**Figure S15**. Box plot of the size distributions of the population of sEVs expressing the three tetraspanins simultaneously for HEK293 and cbMSC cell lines.
